# Global Dynamic Molecular Profiles of Stomatal Lineage Cell Development by Single-Cell RNA Sequencing

**DOI:** 10.1101/2020.02.13.947549

**Authors:** Zhixin Liu, Yaping Zhou, Jinggong Guo, Jiaoai Li, Zixia Tian, Zhinan Zhu, Jiajing Wang, Rui Wu, Bo Zhang, Yongjian Hu, Yijing Sun, Yan Shangguan, Weiqiang Li, Tao Li, Yunhe Hu, Chenxi Guo, Jean-David Rochaix, Yuchen Miao, Xuwu Sun

**Affiliations:** State Key Laboratory of Cotton Biology, Key Laboratory of Plant Stress Biology, School of Life Sciences, Henan University, 85 Minglun Street, Kaifeng 475001, China; College of Life Sciences, Shanghai Normal University, Guilin Road 100, Shanghai, 200234, China; Departments of Molecular Biology and Plant Biology, University of Geneva, Geneva, 1211, Switzerland

## Abstract

The regulation of stomatal lineage cell development has been extensively investigated. However a comprehensive characterization of this biological process based on single-cell transcriptome analysis has not yet been reported. Here, we performed RNA-seq on over 12,844 individual cells from the cotyledons of five-day-old *Arabidopsis seedlings*. We identified 11 cell clusters corresponding mostly to cells at specific stomatal developmental stages with a series of new marker genes. Comparative analysis of genes with the highest variable expression in these cell clusters revealed three transcriptional networks that regulate the development of mesophyll and guard cells, as well as the differentiation from protodermal to guard mother cells. We investigated the developmental dynamics of marker genes via pseudo-time analysis which revealed potential interactions between them. The identification of several novel marker genes suggests new regulatory mechanisms during development of stomatal cell lineage.

## INTRODUCTION

Stomata, which are formed by paired guard cells, have played crucial roles in the colonization of land by plants (von Groll and Altmann, 2001). Turgor-driven stomatal movement requires ion and water exchange with neighboring cells and controls transpiration and gas exchange between plants and the environment. To function efficiently, the development of stomata follows a one-cell-spacing rule, in which two stomata are separated by at least one non-stomatal cell (Bergmann and Sack, 2007; Pillitteri and Torii, 2012). In Arabidopsis, stomata develop from protodermal cells (PDC) through a series of asymmetrical and symmetrical divisions (Han and Torii, 2016). PDCs produce pavement cells (PCs) and self-renewing meristemoids (Ms) that divide asymmetrically several times, generating Ms and PCs known as stomatal lineage ground cells (SLGCs) (Rudall et al., 2013). Ms can eventually differentiate into guard mother cells (GMCs) and a final symmetrical division of a GMC produces two guard cells (GCs) (Geisler et al., 2000). The final spacing between stomata is the result of these M divisions (Pillitteri and Torii, 2012).

Several key genes and regulatory networks underlying stomatal development have been uncovered by molecular and genetic analyses. The closely related basic helix-loop-helix (bHLH) transcription factors SPEECHLESS (SPCH), MUTE, and FAMA control sequential cell fate transitions from meristemoid mother cell (MMC) to M, M to GMC, and GMC to GC, respectively (Ohashi-Ito and Bergmann, 2006; MacAlister et al., 2007; Pillitteri et al., 2007). To specify each cell state differentiation, SPCH, MUTE, and FAMA form heterodimers with two paralogous bHLH-leucine zipper (bHLH-LZ) transcription factors, SCREAM (SCRM) and SCRM2 (Kanaoka et al., 2008). In addition, two partially redundant R2R3 MYB transcription factors, FOUR LIPS (FLP) and MYB88, control stomatal terminal differentiation independently of FAMA (GMC to GCs) (Lai et al., 2005; Ohashi-Ito and Bergmann, 2006). Two secreted cysteine-rich peptides, EPIDERMAL PATTERNING FACTOR1 (*EPF1*) and *EPF2*, are expressed at later and earlier stages of stomatal development, respectively. These peptides are perceived by the cell-surface receptors, ERECTA (ER)-family leucine-rich repeat receptor kinases (LRR-RKs), ER-LIKE1 (*ERL1*) and *ERL2*, resulting in inhibition of stomatal development (Shpak et al., 2005; Hara et al., 2007; Hunt and Gray, 2009b; Lee et al., 2012). The receptor-like protein TOO MANY MOUTHS (TMM) modulates the signaling strength of ER-family receptor kinases in a domain-specific manner (Nadeau and Sack, 2002; Lee et al., 2012). Genetic evidence suggests that these signals are mediated via a mitogen-activated protein kinase (MAPK) cascade, which eventually downregulates the transcription factors responsible for initiating stomatal lineage via direct phosphorylation (Bergmann et al., 2004; Lampard et al., 2008; Lampard et al., 2009; Kim et al., 2012). Stomagen (also known as EPF-LIKE9) peptide promotes stomatal development by competing with EPF2 for binding to ER (Sugano et al., 2010; Zhang et al., 2014; Hronkova et al., 2015). One homeodomain-leucine zipper IV (HD-ZIP IV) protein, HOMEODOMAIN GLABROUS2 (HDG2), acts as a key epidermal component promoting stomatal differentiation (Peterson et al., 2013). It is highly expressed in meristemoids, and a *hdg2* mutant exhibits delayed meristemoid-to-GMC transition (Peterson et al., 2013).

Gene expression profiles for different types of stomatal lineage cells are currently lacking, resulting in a poor understanding of the regulatory mechanisms controlling the PDC to MMC transition. To gain new insights into this process, we isolated protoplasts from cotyledons of five-day-old Arabidopsis seedlings for single-cell RNA sequencing (scRNA-seq). We classified the major cell types and employed transcriptomic analysis to identify several potential key regulators and signaling pathways present in these heterogeneous cell populations. Our analysis led to the identification of a regulatory network of transcription factors for specific developmental stages of stomatal lineage cells. Pseudo-time analysis was employed to uncover the interactions and mutual regulation among key marker genes at different developmental stages. We also identified several novel marker genes that play important roles in regulating stomatal development. These results provide insights into how single-cell transcriptomics can be used to further elucidate the regulatory mechanisms controlling the differentiation of stomatal lineage cells.

## RESULTS

### Gene Expression Pattern of Stomatal Lineage Cells

To systematically resolve gene expression patterns in specific stomatal lineage cells at different developmental stages, we prepared protoplasts by enzymatic digestion from the cotyledons of five-day-old seedlings. The protoplasts were screened with a 40 μm pore cell strainer to obtain more than 20,000 individual cells. Single cells were labeled using 10× Genomics barcode technology, followed by reverse transcription to obtain a single cell cDNA library (Figure S1). This cDNA library was utilized for high throughput sequencing (Figure S1). After extensive analysis of the sequencing results, we obtained transcriptome information for 13,999 single cells (Figure S2). We also identified mitochondrial (mito), chloroplast (pt) and ribosomal (ribo) transcriptomes. Transcripts from these subcellular organelles were excluded from subsequent analysis, resulting in 12,844 single-cell transcriptomes that were further analyzed. They were classified into 11 clusters using t-distributed stochastic neighborhood embedding (t-SNE) (Figure 1). We selected representative marker genes to identify each different cell type: for mesophyll cells (MPC), we used *Ribulose Bisphosphate Carboxylase Small Subunit* (*RBCS*) and *light-harvesting chlorophyll a/b-binding protein* (*LHCB*) as markers that encode chloroplast proteins and are high expressed in MPC; for the epidermal cell populations, we selected *EPF2, BASL, TMM* and *SPCH* as markers for PDC (Pillitteri and Dong, 2013); *POLAR*, *SPCH*, *TMM*, *MUTE*, *HDG2* and *EPF2* were used for MMC (Pillitteri and Dong, 2013); *MUTE*, *BASL*, *SPCH* and *EPF2* were selected for early stage meristemoid (EM) cell identification (Pillitteri and Dong, 2013); *BASL*, *MUTE* and *EPF1* were chosen for late stage meristemoid (LM) cells, while *EPF1*, *HIC*, *FAMA* and *SCRM* were used for GMC (Pillitteri and Dong, 2013); *RBCS*, *FAMA* and *EPF1* were utilized for young guard cells (YGC) (Pillitteri and Dong, 2013); high expression of *HIC*, *RBCS*, *FAMA* combined with low expression of *EPF1* and *EPF2* was used as a marker for GC (Pillitteri and Dong, 2013); *ROP2*, *ROP6*, *ARP2*, *ARP3*, *IQD5* and *RBCS* for PC (Xu et al., 2011; Zhang et al., 2013; Barton et al., 2016; Liang et al., 2018). Because there are chloroplasts in GC, YGC and PC, we also used *RBCS* as marker for these cells (Barton et al., 2016).

**Figure 1.**
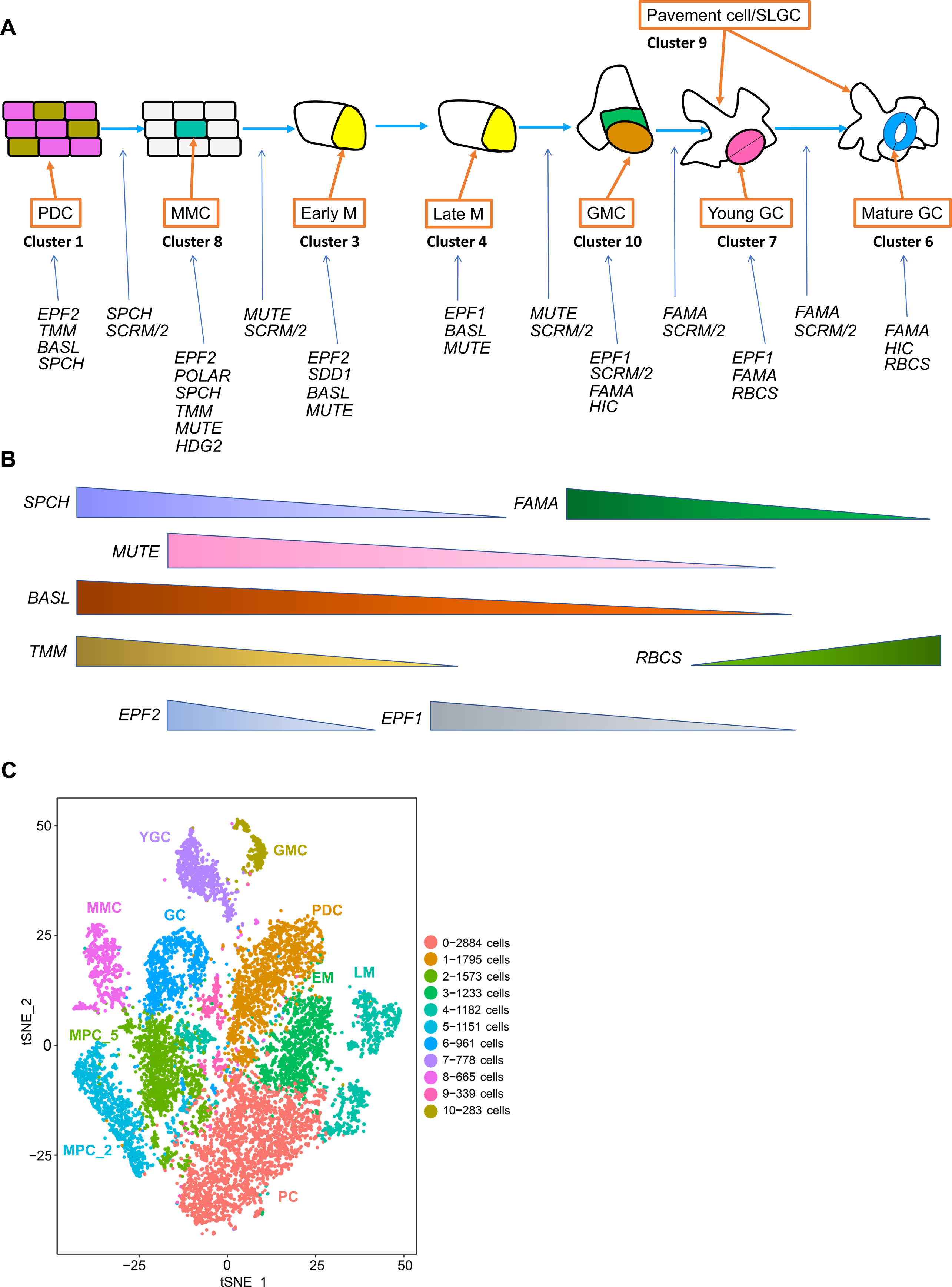
Identification of the cell types with representative marker genes. (**A**) Scheme of expression of marker genes in different cell types. (**B**) Analysis of the dynamic pattern of marker genes during development of stomata. (**C**) Identification of the cell types according to the expression pattern of markers in each cell cluster.

The expression profiles of the above selected marker genes have been determined previously. *SPCH* is expressed in the developing leaf epidermis (MacAlister et al., 2007). A transcriptional green fluorescent protein (GFP) reporter (*SPCHpro::nucGFP*) and a translational reporter (*SPCHpro::SPCH-GFP*) are expressed in a subset of epidermal cells that lack overt signs of differentiation (MacAlister et al., 2007). In cotyledons, *SPCH* expression is often observed in two neighbouring cells, a pattern consistent with expression in the dividing cell population (MacAlister et al., 2007). In older organs, *SPCHpro::SPCH-GFP* expression continues to be restricted to small cells in the epidermis, including cells that have recently divided next to stomatal lineage cells (MacAlister et al., 2007). In meristemoids, *SPCH* expression is downregulated and *MUTE* expression commences (MacAlister et al., 2007). *MUTE* is required to limit the number of rounds of meristemoid division and expressed strongly in meristemoids and at lower levels in GMCs and GCs (Pillitteri et al., 2007). *FAMA* is expressed in GMCs and is necessary and sufficient to promote GC identity (Ohashi-Ito and Bergmann, 2006). The *PROFAMA:GFP* expression is restricted to the stomatal lineage (Ohashi-Ito and Bergmann, 2006). The *PROFAMA:GFP* is not expressed in meristemoids but is strongly expressed in GMCs and in YGCs (Ohashi-Ito and Bergmann, 2006). BASL and POLAR show largely overlapping localization at the cell cortex during stomatal asymmetric divisions (Dong et al., 2009). In the cotyledon and leaf epidermis, *BASL::GUS* is highly expressed in the asymmetrically dividing MMCs and meristemoids, and it is undetectable in later stomatal lineage cells (Dong et al., 2009). *ProTMM::TMM-GFP* is expressed in proliferating stomatal lineage cells, but not in other epidermal cells or in mature stomata (Nadeau and Sack, 2002). EPF2 is produced in *SPCH*-expressing PDC (MMCs) early in the lineage, whereas EPF1 is produced in late-stage meristemoids, GMCs and young guard cells (Hara et al., 2007; Hunt and Gray, 2009a). Consistent with the report on *HDG2pro::GUS* (Nakamura et al., 2006), the GFP signals of the HDG2 transcriptional reporter (*HDG2pro::nls-3xGFP*) are strongly expressed in meristemoids and SLGCs (Peterson et al., 2013). The HDG2 translational reporter (*HDG2pro::HDG2-GFP*) accumulates in the nuclei of meristemoids and SLGCs (Peterson et al., 2013). ROP2 and ROP6 are locally activated at the opposing sides of the cell wall and are mutually exclusive along the plasma membrane within PCs (Fu et al., 2005; Xu et al., 2011). In addition, microtubule-associated protein IQ67 DOMAIN 5 (IQD5) is localized in PC (Liang et al., 2018). The *GFP:IQD5* colocalizes with the microtubule marker *mCherry:TUB6* (mCherry fused to β-tubulin6) in both leaf PC and hypocotyl epidermal cells (Liang et al., 2018). The scheme in Figure 1A, B displays the expression pattern of selected marker genes in different stomatal cell types.

### Identification of the cell types with marker genes

To determine the cell type with the above maker genes, we analyzed the pattern of selected marker genes in each cell cluster. As shown in Figure S3A and B, *FAMA* expression is high in clusters 6 and 7. *SCRM* is expressed in clusters 6,7,8 and 10 (Figure S3A and B). *SPCH* expression is high in clusters 7,8 and 9 (Figure S3A). *MUTE* is expressed in cluster 8 (Figure S3A). *BASL* is expressed in clusters 1, 4 and 10 (Figure S3A). High expression of *POLAR* is found in clusters 6, 7 and 8 (Figure S3A). *EPF1* is expressed in clusters 6, 7 and 8, while *EPF2* is expressed in clusters 6, 7, 9 and 10 (Figure S3A). *EPFL9*, *ROP2*, *ROP6* and *IQD5* are highly expressed in cluster 0 (Figure S3A). Based on the expression patterns of these marker genes, we can determine the cell type of each cluster as follows: cluster 0 is PC, cluster 1 is PDC, cluster 8 is MMC, cluster 3 is EM, cluster 4 is LM, cluster 10 is GMC, cluster 7 is YGC and cluster 6 is GC (Figure 1C). The expression of *RBCS* and *PSAB* is mainly enriched in clusters 2 and 5 indicating that they correspond to MPCs. To distinguish them, we named them MPC_2 and MPC_5 respectively (Figure 1C). For cluster 9, we could not determine its cell type with the known marker genes. However amongst the genes that belong to this cluster, we checked the stomatal pattern in the corresponding mutants. As an example we found that *BCL-2-ASSOCIATED ATHANOGENE 6* (*bag6*) affects the distribution of GC and produces some double and adjacent GCs, as well as an increase of SI compared with WT (Figure 1A, Figure S4). Plant BAG proteins are homologs of mammalian regulators of apoptosis and play important roles in regulating apoptotic-like processes ranging from pathogen attack, to abiotic stress, to plant development (Kabbage and Dickman, 2008; Li et al., 2016; Fu et al., 2019). Expression of *BAG6* in leaves was strongly induced by heat stress (Fu et al., 2019). Loss of function of BAG6 in *fes1a bag6* double mutant partially rescued the sensitive of *fes1a* to heat stress (Fu et al., 2019). The *bag6* mutant shows enhanced susceptibility to the fungal pathogen *Botrytis cinerea*, both in terms of severity of lesions and rate of spread (Li et al., 2016). Since stomata are important entry sites for fungal pathogens (Melotto et al., 2006; Underwood et al., 2007; Dutton et al., 2019; Zhang et al., 2019), the increased number of stomata in *bag6* may lead to the fast spread of *Botrytis cinerea*. Another mutant from cluster 9 marker genes, *bzip6*, does not significantly affect the development of stomata. The expression of *bZIP6::ER-GFP* occurs specifically in two pericycles in the phloem pole starting from the early root elongation zone (Lee et al., 2006) . These results suggest that *bZIP6* is not a marker gene of stomata. and that cluster 9 does not belong to epidermal cells although the *bag6* mutant is defective in the distribution of GC.

To investigate the abundance of gene transcripts in different cell types, we counted the number of cells and the number of transcripts identified in each cell type (Figure S5A and B and Supplemental Table S1). Note that the number of different cell types identified here does not directly reflect the relative number of different cell types in the cotyledons of plants. The number of identified cells only reflects the relative number of each cell type in the cell samples we obtained. At the same time, the number of transcripts identified in each cell type was also quantitatively analyzed by determining the average number of transcripts identified in each cell for comparison (Figure S5C). The results show that the number of average transcripts in PC was lowest, whereas it was highest in GMC. In contrast, a relatively high number of transcripts was identified in MMC, PDC and EM (Figure S5C).

### Expression of marker genes in stomatal lineage cells

To further test the cell type that we identified, we analyzed the expression of several known marker genes that are involved in regulating the development of stomatal lineage cells. As shown in Figure S3B, *FAMA*, *TMM*, *HIC* and *SCRM* are specifically expressed in YGC and GMC, while other marker genes are not only expressed in YGC and GMC, but also in other cell types (Figure S3B), suggesting that their functions may not be only restricted to the regulation of stomatal lineage cell development. To explore the potential regulators of stomatal lineage cells, we analyzed gene expression profiles in different clusters and identified highly expressed marker genes in each individual cell cluster (Figure 2A). Feature plot analysis indicated that the expression of newly identified marker genes is clearly increased in their corresponding clusters (Figure 2B and Supplemental Table S2). Some of these marker genes could potentially be involved in regulating the development of stomatal lineage cells. SLAC1 and SCAP1(DOF5.7) play important roles in regulating the development of stomatal lineage cells (Engineer et al., 2015; Chen et al., 2016). Consistently, a high level of expression of *SLAC1* and *DOF5.7* was detected in YGC and GMC (Figure 2A). Furthermore, a pectin methylesterase gene, *PME6*, is highly expressed in YGC and GMC (Figure 2A). As reported, *PME6* is highly expressed in guard cells and required for stomatal function (Amsbury et al., 2016). Guard cells from *pme6-1* mutant have walls enriched in methyl-esterified pectin and show a decreased dynamic range in response to elevated osmoticum, suggesting that the mechanical change in the guard cell wall can affect stomatal function (Amsbury et al., 2016). The Arabidopsis K^+^ channel gene, *KAT2*, is expressed in guard cells (Pilot et al., 2001). KAT2 is a major determinant of the inward K^+^ current through the guard cell membrane (Pilot et al., 2001). Interestingly, the expression of *ARABIDOPSIS THALIANA MERISTEM LAYER1*(*ATML1*) was high in MMC, where the established marker genes *SPCH* and *MUTE* were also highly expressed (Figure 2A).

**Figure 2.**
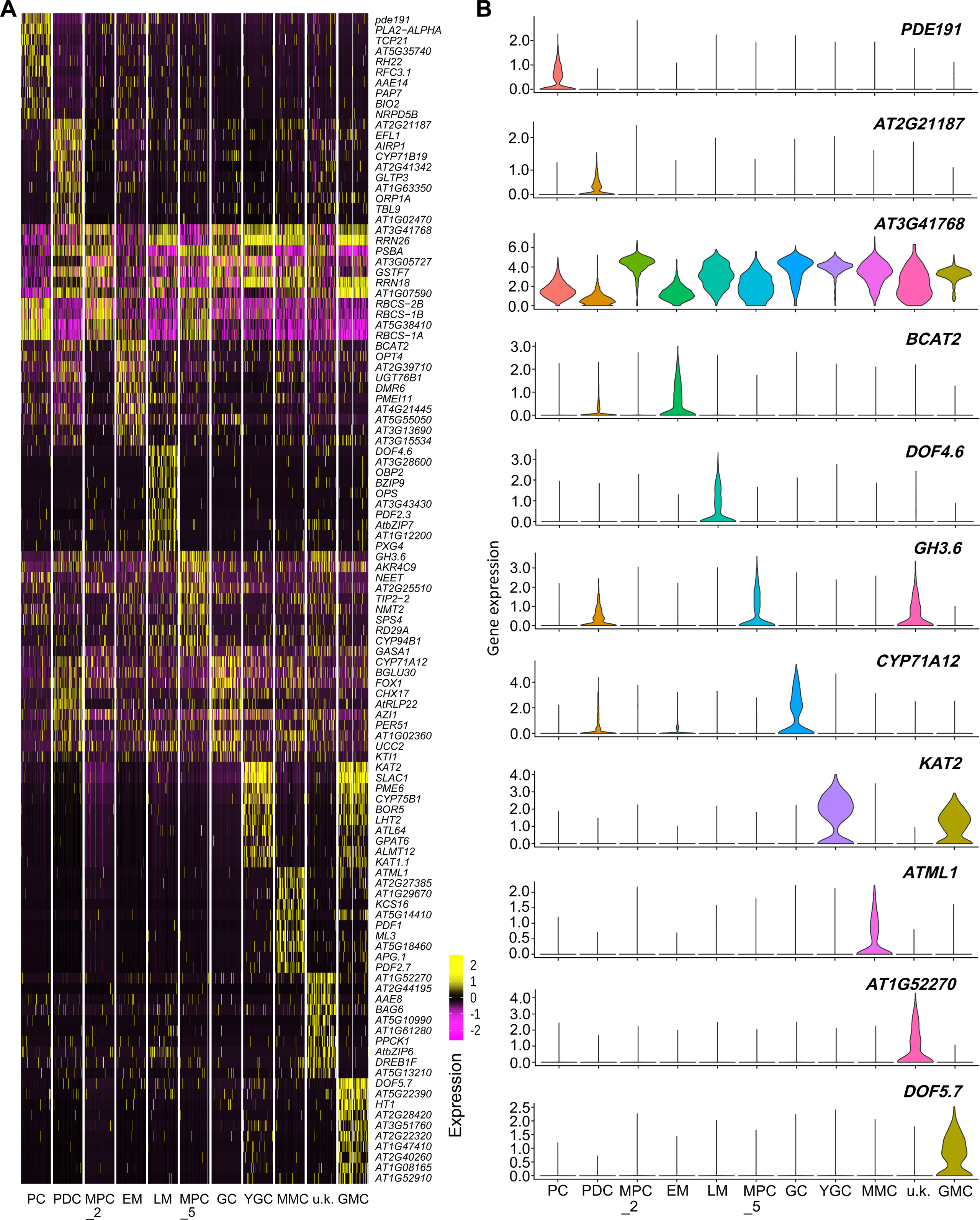
Identification of novel marker genes for each cluster. (**A**) Heatmap of expression of representative marker genes in each cluster. (**B**) Violin_plots show expression of representative marker genes in each of cell types. PDC: protodermal cells, PC: pavement cell, M: meristemoid, GMC: guard mother cell, GC: guard cell, MMC: meristemoid mother cell, EM: early stage meristemoid, LM: late stage meristemoid, YGC: young guard cell, MPC: mesophyll cell, u.k.: unknown.

In GC, we identified 10 top marker genes that may be involved in regulating the function of GC (Figure 2A). Stomata are not only required for gas exchange with the environment and for controlling water loss, but they also provide routes for pathogen entry into plants (Zeng et al., 2010). Therefore plants have evolved mechanisms to regulate stomatal aperture as an immune response against bacterial invasion (Zeng et al., 2010). A recent study showed that bacterial infection can induce systemic signaling to inhibit the development of stomata in new leaves to restrict pathogen entry (Dutton et al., 2019). The bacterial peptide flg22 or the phytohormone salicylic acid (SA) induce a similar systemic decrease in stomatal density (Dutton et al., 2019). This developmental process can be regulated by AZELAIC ACID INDUCED 1 (AZI1), which is involved in the priming of SA induction and systemic immunity triggered by pathogen or azelaic acid (Pitzschke et al., 2016). *KTI1* encodes an Arabidopsis serine protease (Kunitz trypsin) inhibitor and plays important roles in modulating PCD in plant-pathogen interactions (Li et al., 2008). Expression of *AtKTI1* is induced late in response to bacterial and fungal elicitors and to salicylic acid (Li et al., 2008). Besides SA, camalexin is also involved in regulating the defense response of plants. One of cytochrome P450 enzymes, CYP71A12 is an important component in the biosynthetic pathway of camalexin and related metabolites (Mucha et al., 2019). Pathogen infection can induce the expression of *CYP71A12*, which leads to dehydration of IAOx to form indole-3-acetonitrile (IAN) during the biosynthesis of camalexin (Nafisi et al., 2007; Muller et al., 2015). CYP71A12 is involved in the biosynthetic pathway to 4-hydroxyindole-3-carbonyl nitrile (4-OH-ICN), the level of all ICN derivatives with the exception of A6 in *cyp71a12* mutant is about 10% of WT (Rajniak et al., 2015). 4-OH-ICN plays roles in inducing the pathogen defense (Rajniak et al., 2015). *CYP71A12* is co-expressed with a flavin-dependent oxidoreductase 1 (*FOX1*), and levels of ICN metabolites in the *fox1* mutant are decreased three-to fivefold compared with WT (Rajniak et al., 2015).

Consistent with these findings, *KT1*, *CYP71A12* and *FOX1* are expressed in GC (Figure 2A) further supporting that stomatal cells may fight against pathogens by producing IAN and OCN. Opening of stomata in response to various stimuli is regulated by K^+^ uptake through inward-rectifying K^+^ channels in the plasma membrane (Szyroki et al., 2001). *CHX17* encodes a putative K^+^/H^+^ exchanger.

*CHX17* cDNA complements the phenotypes of the *kha1Delta* mutation in S. cerevisiae cells, which shows a growth defect at increased pH and hygromycin sensitivity (Maresova and Sychrova, 2006). Under its native promoter, AtCHX17(1-820)-GFP is localized in the prevacuolar compartment and in plasma membrane in roots (Chanroj et al., 2013). Expression of *CHX17* in GC suggests that it may be involved in regulating the opening of stomata.

### ATML1 is involved in regulating the development of stomatal lineage cells

To explore the potential regulator of MMC, we analyzed the marker genes in MMC and found that *ATML1*, *PDF1*, *MUTE* and *SPCH* have highly similar expression profiles (Figure 2A and Figure S3A). To investigate the roles of ATML1 in the regulation of epidermal cell differentiation, Takada et al generated a construct *proRPS5A-ATML1* that uses the promoter region of *AtRPS5A* to drive the expression of *ATML1* (Takada et al., 2013). The resulting construct was transformed into a transgenic plant containing *STOMAGEN-GUS* to investigate the effects of ATML1 on the expression of the mesophyll-specific *STOMAGEN-GUS* reporter (Takada et al., 2013). In the transgenic plants of *proRPS5A-ATML1*, the expression of *ATML1* was induced by treating the seedlings with β-estradiol (Takada et al., 2013). Overexpression of *ATML1* induced stomata-like structures in the inner cells of the cotyledons in independent lines (Takada et al., 2013). These ectopic guard cell-like cells expressed the guard cell marker *KAT1-GUS*, suggesting that these cells have guard cell identity (Takada et al., 2013). This result also suggested that overexpression of *ATML1* can induce the development of stomata. Moreover, induction of *ATML1* can inhibit the expression of mesophyll-specific *STOMAGEN-GUS* and result in miss-shaped leaves with ectopic patches of transparent cells among the green mesophyll tissues (Takada et al., 2013). These results suggest that induction of *ATML1* can enhance the biogenesis of stomata but inhibit the development of mesophyll tissues. Although STOMAGEN can enhance the biogenesis of stomata (Lee et al., 2015), induction of *ATML1* does not inhibit the biogenesis of stomata even if the expression of *STOMAGEN* is suppressed by ATML1, suggesting that ATML1 may rely on a STOMAGEN-independent pathway to enhance the biogenesis of stomata. Furthermore, the SI of the *atml* mutant is decreased, while the SI of ATML1-OX transgenic plant overexpressing *ATML1* is increased (Peterson et al., 2013), suggesting that ATML1 is involved in regulating the development of stomata. To investigate the effects of ATML1 on the biogenesis of stomatal lineage cells, we used mutants of ATML1. As expected, *atml1-2* and *atml1-3* plants are deficient in the development of stomatal lineage cells (Figure 3A-F). Further RT-PCR analysis indicated that the expression levels of *SPCH* and *MUTE* in *atml1-2* and *atml1-3* were lower than in WT (Figure 3G), suggesting that ATML1 can regulate the development of stomatal lineage cells by modulating the expression of both *SPCH* and *MUTE*.

**Figure 3.**
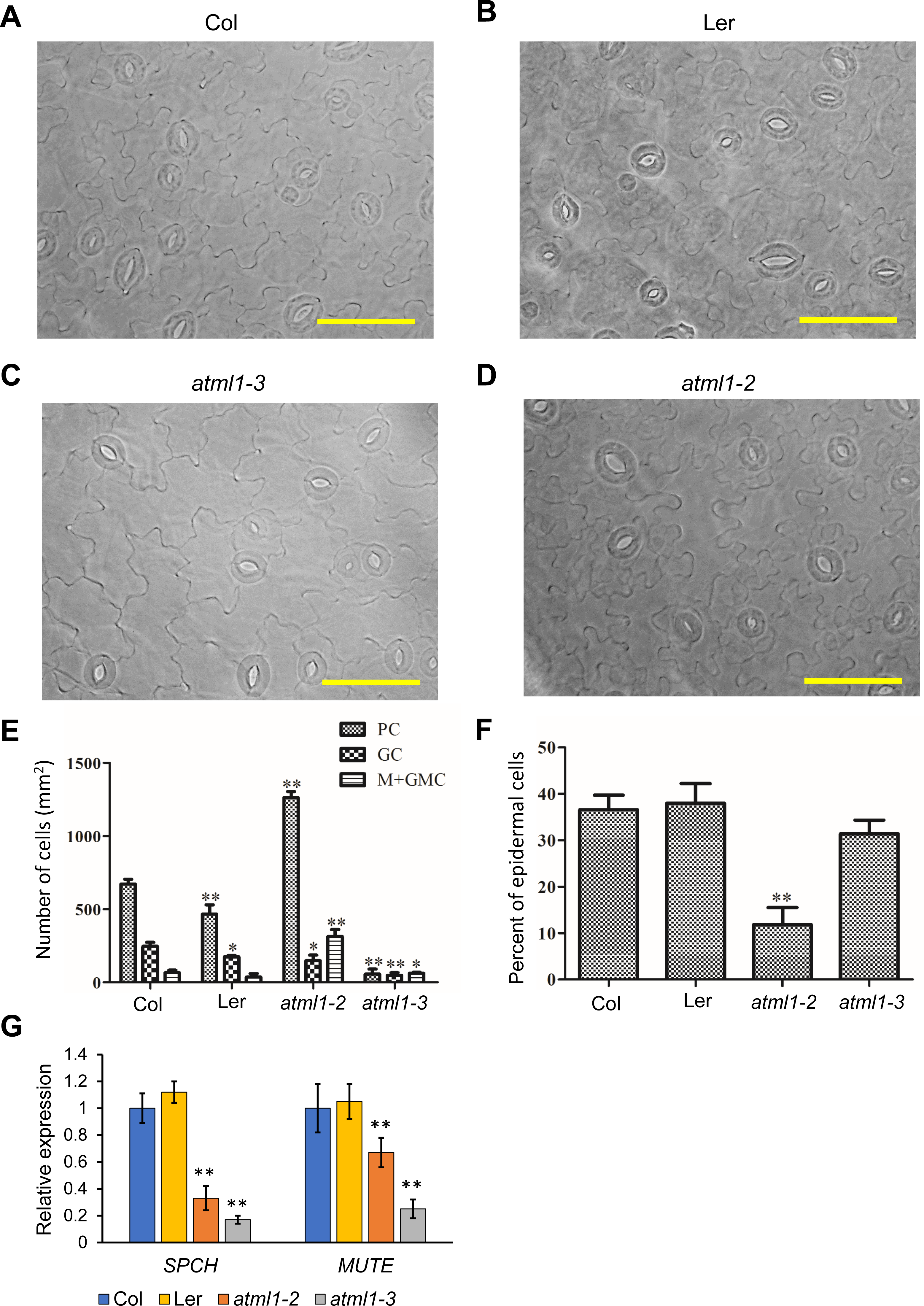
ATML1 is involved in regulating the development of stomata. (**A**-**D**) Analysis of stomatal development of 5-day-old seedlings of *atml1-2* (in Ler background), *atml1-3* (in Col background), Ler and Col are used as controls. (**E-F**) Quantitative analysis of **A**-**D**. (**G**) qPCR analysis of the expression of *SPCH* and *MUTE* in 5-day-old seedlings of *atml1-2*, *atml1-3*, Ler and Col. Error bars represent standard errors (S.E.). *: p<0.05, **: p<0.01, one-way ANOVA analysis versus Col. Scale bar: 50 μm in **A**-**D**.

### GO Analysis of the genes enriched in different cell types

To investigate the potential biological function of genes expressed in each cell type, we performed GO analysis on all cell clusters (Figure 4 and Figure S6). There were significant differences in the number of enriched genes identified in different cell types. In general, the majority of enriched terms were associated with individual cell types, however those GO terms associated with multiple cell types represent more general biological processes (e.g., response to oxidative stress and salt stress, and vesicle-mediated transport) (Figure 4A and Figure S6). As a measure of the reliability of our method in identifying cell type–expressed genes and of our ability to correctly annotate biological processes to a cell type, we compared a list of genes enriched in GCs in our analysis with a previous study that profiled GC functions. In agreement with these published reports (Gray, 2005; Lawson, 2009; Song et al., 2014; Niu et al., 2018; Huang et al., 2019), we found genes that respond to oxidative stress, salt stress, bacteria, cadmium ions and are involved in stomatal movement and photosynthesis, are preferentially expressed in GCs (Figure 4A and Figure S6). Our analysis further increased the spectrum of biological processes associated with GC development to include protein transport, vesicle-mediated transport, and cell death (Figure 4A and Figure S6). Since this result indicated that our method is suitable, we used gene categories that are preferentially expressed to infer the function of other cell types. Gene ontology (GO) heatmap analysis indicated that the genes expressed in PC and MPC are mainly involved in photosynthesis and carbohydrate metabolism (Figure 4A). The genes with increased expression in GCs and YGCs are also involved in photosynthesis, which is consistent with the presence of chloroplasts in these cells (Figure 4A). The genes expressed in cluster 9 are mainly implicated in the response to abiotic and biotic stress and protein transport (Figure 4A). We could not assign a cell type to cluster 9 due to the uncertainty about marker genes (Figure 4A). GO heatmap analysis revealed that MMC, EM, LM and GMC are similar (Figure 4A and B). In these cells, the expressed genes are not involved in photosynthesis (Figure 4A). Unexpectedly, we found that genes preferentially expressed in these cells are involved in regulating the response to all kinds of environmental stress and stimulus (Figure 4A and B and Figure S6). For instance, genes that respond to bacteria are highly expressed in MMC (Figure 4B). Compared with LM and GMC, the most important feature of MMC is the lack of gene expression associated with the respiratory chain and oxidative phosphorylation (Figure 4B), suggesting that MMCs have relatively low metabolic activity. However, genes enriched in PDC and MMC are involved in protein transport, vesicle-mediated transport and membrane protein complexes (Figure 4A and Figure S6), suggesting that PDC and MMC show higher activity of protein expression.

**Figure 4.**
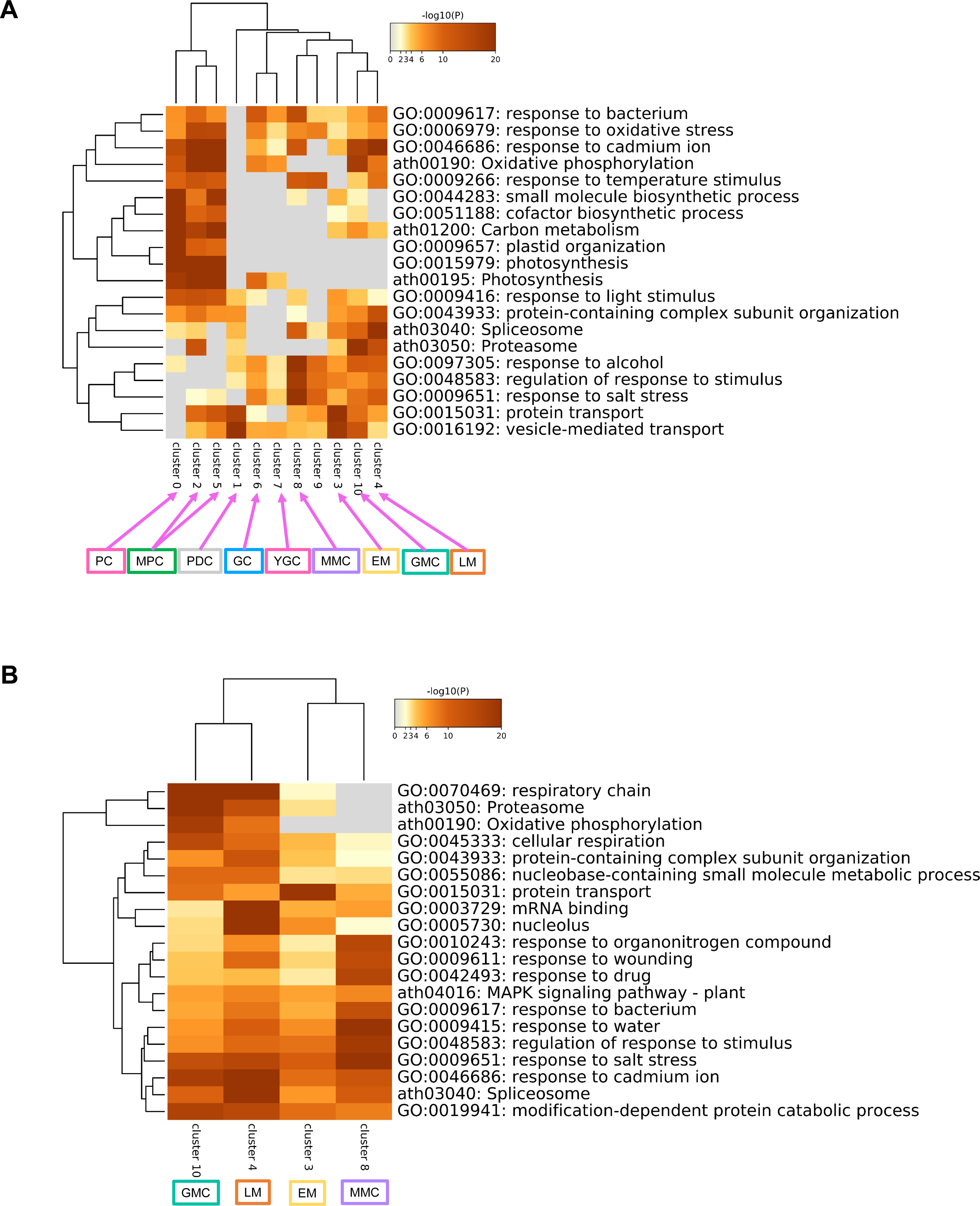
GO analysis of the genes that expressed in different cell types. (**A**) GO-heatmap analysis of the genes with the highest variable expression in different clusters. (**B**) Same analysis as in A for MMC, GMC, EM and LM.

### Analysis of the regulatory network of transcription factors (TFs) in different cell types

To investigate the mechanisms that regulate the development of different cell types, we analyzed the regulatory network of TFs in each of them. We first analyzed the number of TFs in different cell types, as shown in Figure 5A. In PDC, we identified the highest number of TFs, while in MPC_5, we identified the lowest number of TFs. More TFs were identified in EM, LM, MMC and GMC, but less in GC and YGC.

**Figure 5.**
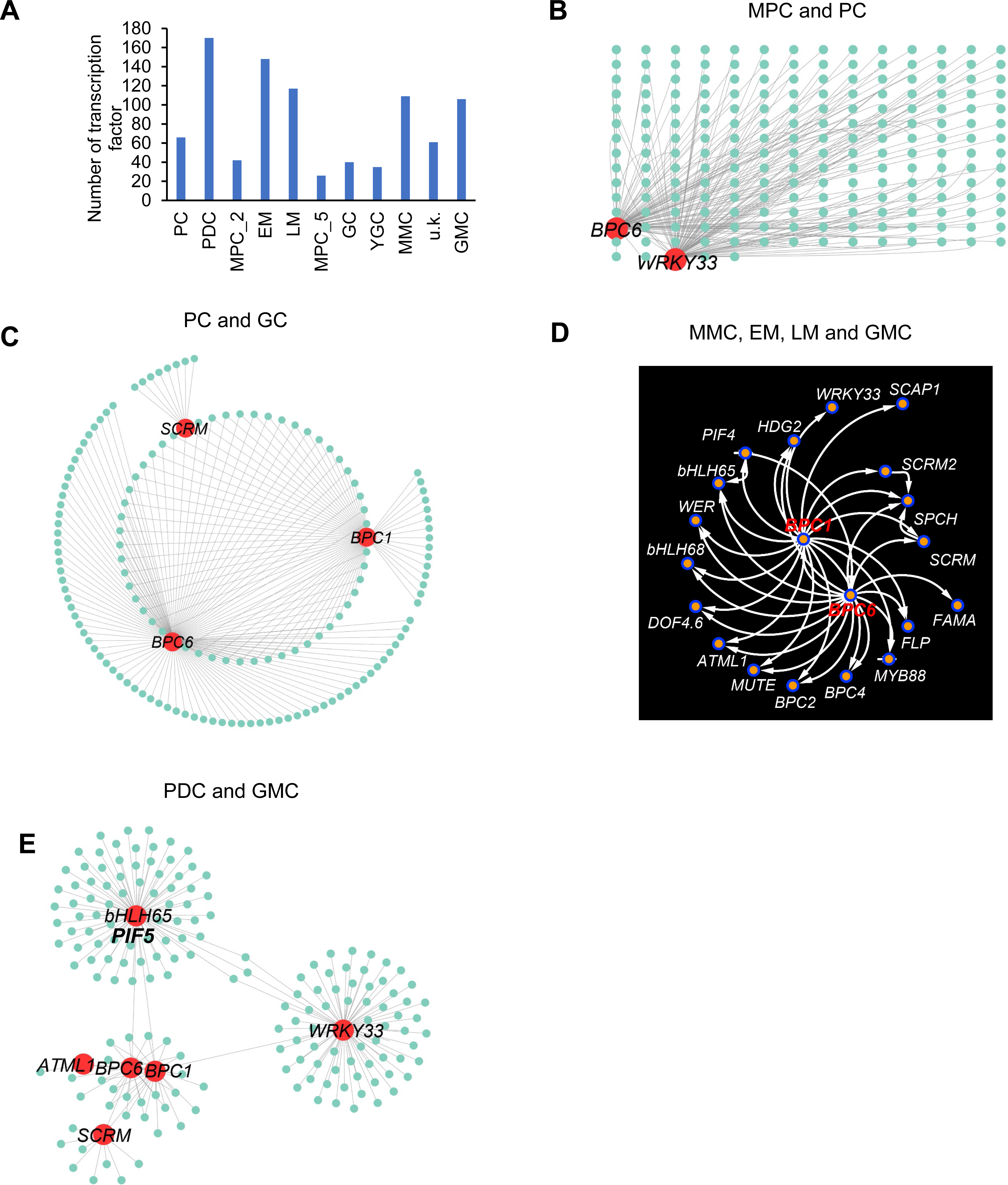
Identification of regulatory networks of transcription factors in different cell types. (**A**) Identification of the transcription factors in different cell types. (**B**) Analysis of the regulatory network of transcription factors in MPC and PC. (**C**) Analysis of the regulatory network of transcription factors in PC and GC. (**D**) Analysis of the regulatory network of transcription factors in MMC, EM, LM and GMC. (**E**) Analysis of the regulatory network of transcription factors in PDC and GMC.

Accumulation of TFs in PDC, EM, LM, MMC and GMC showed that gene expression was higher at the early stage of development of stomatal cells, and the number of TFs needed was also higher. In GC and YGC, gene expression was relatively low. Surprisingly, we found that the number of TFs in MPC_2 and MPC_5 was lower than in other cell types. Although MPCs are important for photosynthesis and for a series of important metabolic reactions, their gene expression is relatively low. This is consistent with the low average number of highly expressed genes in MPC cells (Figure S5A and B). The mRNAs of the transcription factors (TFs) *BASIC PENTACYSTEINE1*(*BPC1*), *BPC6*, and *WRKY33* are highly expressed in PC and MPC (Figure 5A). We further analyzed the regulatory network of these TFs by analyzing the genes co-expressed with them and extracted the top 1,000 links showing positive correlation with BPC6 and WRKY33 (Figure 5A). We found that BPC6 and WRKY33 are core TFs in regulating the development and function of PC and MPC (Figure 5A). Analysis of the regulatory network of TFs in YGC and GC suggests that BPC1, BPC6 and SCRM may act as the core TFs regulating the development and function of PC and GC (Figure 5B). SCRM has been shown to interact with FAMA to regulate the differentiation from GMC to GC (Kanaoka et al., 2008). Thus, our results suggest that BPC1 and BPC6 may mediate the development of GC in conjunction with SCRM. Analysis of the regulatory network of TFs in MMC, EM, LM and GMC indicates that they have fewer close interactions with known transcription factors (except for SCRM, SPCH and SCRM2), but they are all associated with *BPC1* and *BPC6* based on co-expression (Figure 5C). There is also a close regulatory relationship between BPC1 and BPC6, which form the core of the transcriptional regulatory network in these cell types (Figure 5C). This finding suggests that BPC1 and BPC6 may regulate the differentiation from PDC to GMC by interacting with other transcription factors. Furthermore, we also found that PIF5, BPC1, BPC6, WRKY33, ATML1, and SCRM can act as core TFs to regulate the differentiation from PDC to GMC (Figure 5D). Feature plot analysis indicated that expression of *BPC1*, *BPC2*, *BPC4* and *BPC6* is mainly enhanced in MMC, EM, LM, GMC, YGC and GC, while the expression of *WRKY33* can be detected in all cell types (Figure 6A). Analysis of the stomatal developmental pattern indicated that the number of GC in *wrky33* is decreased, while the numbers of M and GMC in *wrky33* are increased, compared with WT (Figure 6B and C), suggesting that WRKY33 is involved in regulating the development of GC from GMC.

**Figure 6.**
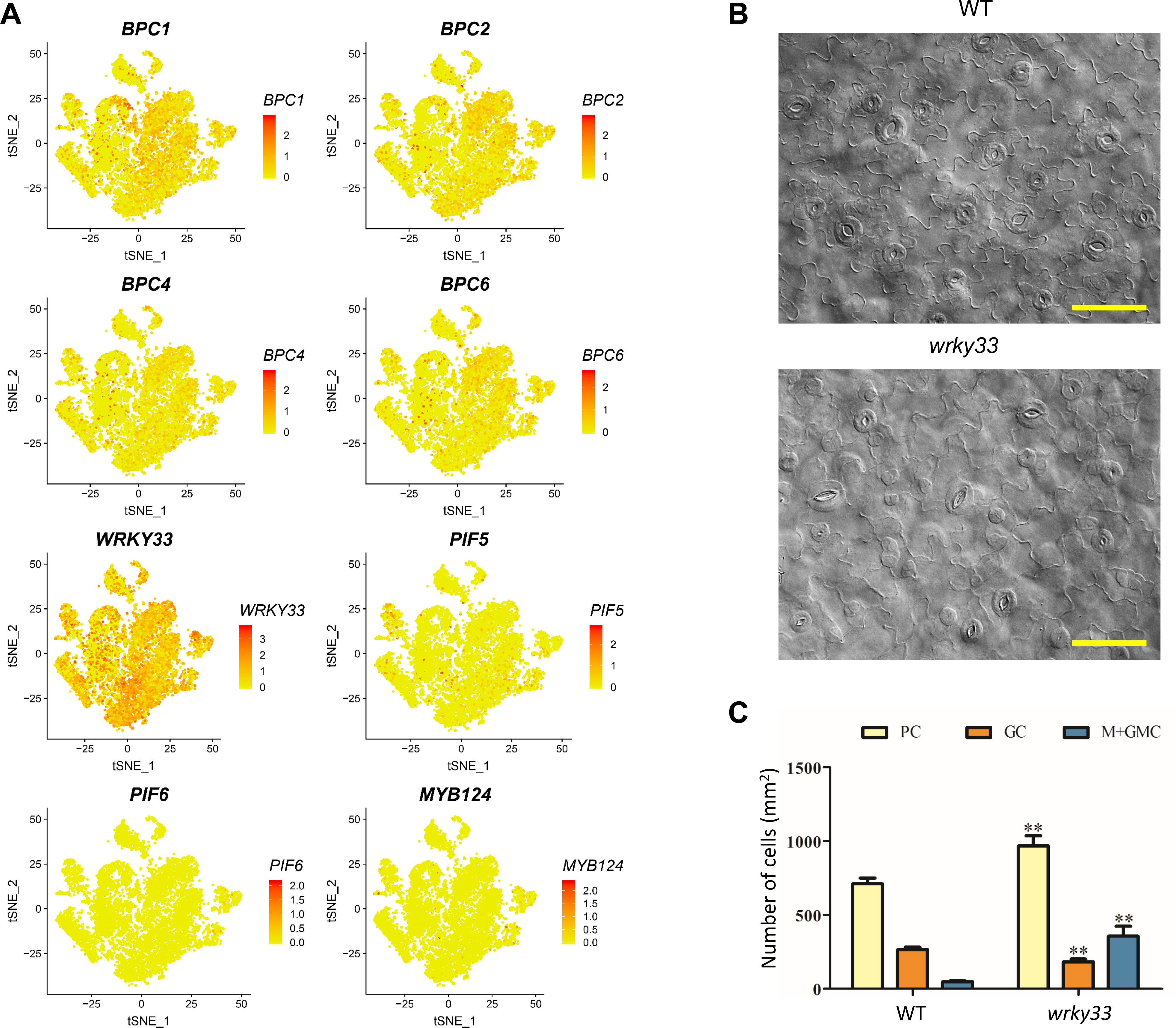
Analysis of feature plots and function of core TFs. (**A**) Feature plots of the expression of representative TFs in different clusters. (**B**) Developmental pattern of stomatal lineage cells of cotyledons of five-day-old seedlings of *wrky33*, with wild type (WT) used as control. (**C**) Frequency of cell types calculated from (**B**). Error bars represent standard errors (S.E.). *: p<0.05, **: p<0.01, one-way ANOVA analysis versus WT. Scale bar: 50 μm in **B.**

BPC6 has been shown to participate in the regulation of *ABI4* (Mu et al., 2017), and subcellular localization indicated that BPC6-GFP is in the nucleus of guard cells (Figure 7A). To detect whether BPCs can directly regulate the expression of key marker genes of stomatal development, we analyzed the transcript levels of both *SCRM* and *SCRM2*, which can form a complex to regulate the functions of SPCH, MUTE, and FAMA (Pillitteri and Torii, 2012). RT-PCR analysis indicated that the expression levels of *SCRM* and *SCRM2* in the *bpc1 bpc2 bpc3 bpc4 bpc6 bpc7* sextuple mutant are lower than in WT (Figure 7B). Further analysis showed that the Stomatal Index (SI) was increased in the *bpc1 bpc2 bpc4 bpc6* quadruple mutant and the *bpc1 bpc2 bpc3 bpc4 bpc6 bpc7* sextuple mutant, whereas the SI of BPC6-GFP was decreased compared with WT (Figure 7C-E). These results suggest that BPCs can mediate stomatal development by regulating the expression of *SCRM* and *SCRM2*.

**Figure 7.**
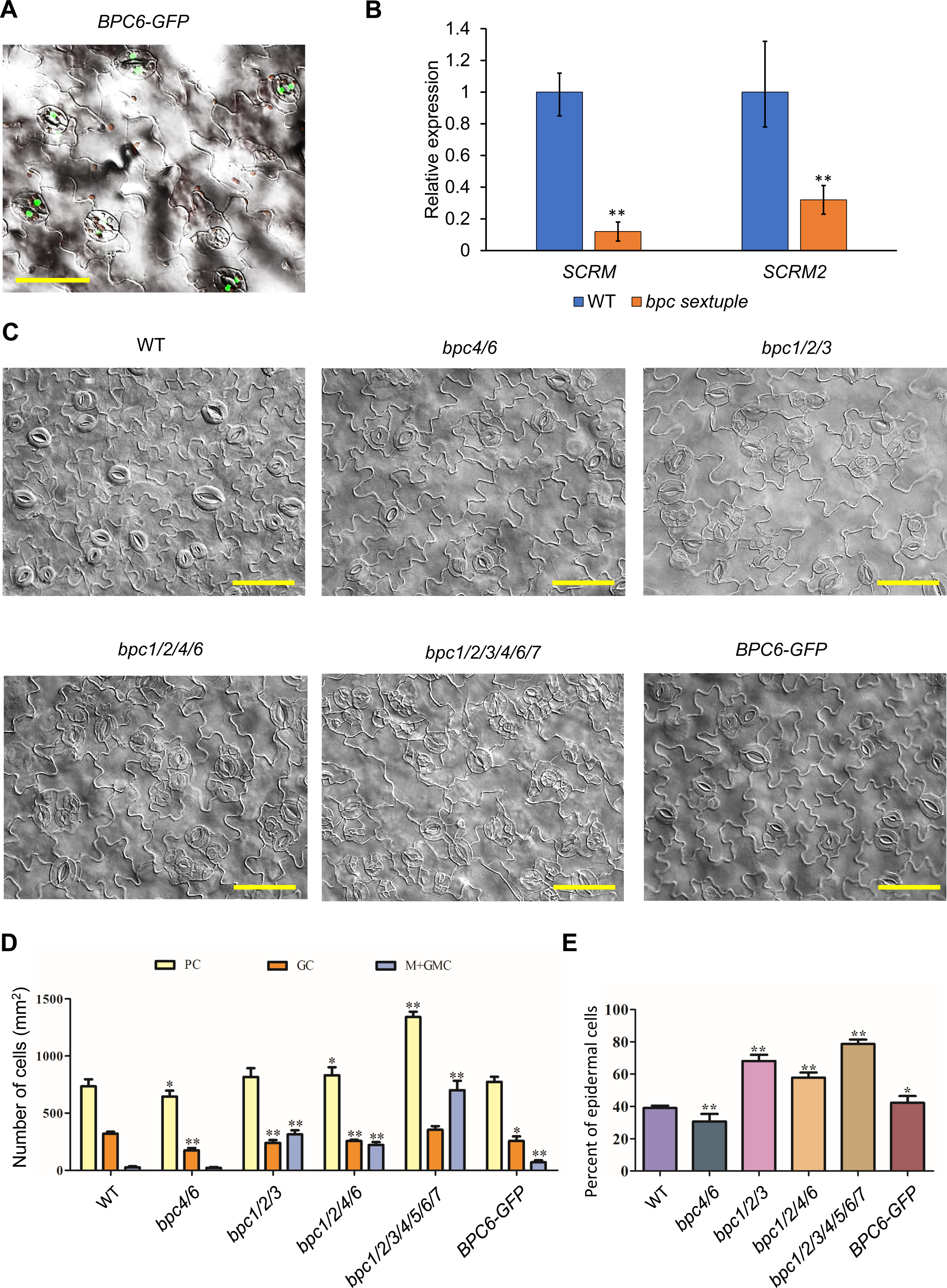
BPC proteins are involved in regulating the development of stomata. (**A**) Analysis of the subcellular localization of BPC6-GFP. (**B**) qPCR analysis of the expression of *SCRM* and *SCRM2* in the *bpc* sextuple mutant. (**C**) Developmental patterns of stomatal lineage cells in cotyledons of 5-day-old seedlings of *bpc* mutants and transgenic plants, WT was use as control. (**D**) Frequency of cell types calculated from (**C**). (**E**) Number of epidermal cells of mutants grown on MS medium. Error bars represent standard errors (S.E.). *: p<0.05, **: p<0.01, one-way ANOVA analysis versus WT. Scale bar: 50 μm in **A** and **C**.

### Developmental Pseudo-time Analysis of Marker Gene Expression

To reconstruct the developmental trajectory during differentiation, we performed pseudo-temporal ordering of cells (pseudo-time) from our scRNAseq data using Monocle 2(Trapnell et al., 2014). In total, the pseudo-time path has three branches (Figure 8A and B), and different cell clusters can be arranged relatively clearly at different branch sites of the pseudo-time path (Figure 8B). In general, the different developmental processes of stomatal lineage cells can be seen from PDC to GC (Figure 8B). Surprisingly, PC was concentrated in a pseudo-time branch that was significantly different from the other cell types (Figure 8B). Intriguingly, we found that PDC and GMC could not be clearly distinguished on the pseudo-time curve (Figure 8B). In principle, the distribution characteristics of different cell types on the development trajectory can preliminarily determine the relationship between these cells during the development period. The distribution of GMC and YGC in the development trajectory is relatively concentrated, but MPC_2 and MPC_5 can be found at several time points along the development trajectory, suggesting that the cell development stage of MPC is more complex. Interestingly, we found that PDC and EM show relatively similar distribution patterns in the development trajectory, suggesting that their developmental stages are close. Although PC is mainly distributed in branch 1 at the end stage of the development trajectory, it is also distributed at other time points, which is consistent with the earlier development stage of PC cells compared to stomatal cells. To investigate the pseudo-time patterns of genes in each cluster, we performed heatmap analysis for all the highly expressed genes (Figure 8C). In general, the pseudo-time patterns of all genes can be divided into three clusters (Figure 8C). To analyze the pseudo-time patterns of representative marker genes, we selected the top 5 marker genes in each cluster to analyze their pseudo-time patterns (Figure 8D). As shown in Figure 8D, the heatmap of pseudo-time of the top 5 marker genes shows that their pseudo-time pattern can be classified into two clusters (Figure 8D). In the first cluster, the expression of all the marker genes increases gradually along with the pseudo-time (Figure 8D). In contrast, expression of marker genes in cluster 2 decreases at the end of the pseudo-time axis (Figure 8D). In the first cluster, the marker genes are mainly from PC, suggesting that PCs are at a more mature stage of development (Figure 8D). The second cluster can be divided into three branches: in the first, expression of marker genes mainly from GMC,YGC,GC and MPC first gradually increases to a maximum level, and then decreases quickly along the pseudo-time; in the second branch, expression of marker genes mainly from PDC and LM is very high at the beginning of development, but quickly decreases along the pseudo-time; in the third branch, expression of the marker genes mainly from MMC and EM is lower in all of the developmental periods along the pseudo-time, and further declines at the last stage of pseudo-time (Figure 8D). It can be seen from the heatmap and curves of pseudo-time that expression of *SPCH*, *MUTE* and *FAMA* occurs mainly between PDC and MMC, MMC and M, and GMC and GC (Figure 9A and B). Expression of the marker genes *EPF1*, *SPCH* and *MUTE* is highly similar at the early stages of development (Figure 9A and B). As expected, *EPF2*, *FAMA* and *SCRM* are mainly expressed in the EM and LM stages and exhibit a similar pseudo-time pattern (Figure 9A and B). Although *SCRM* and *SCRM2* have some functional interactions with *MUTE* and *SPCH* in the regulation of stomatal lineage cell development, our pseudo-time results show that expression of the *SCRM*, *SCRM2*, *MUTE* and *SPCH* genes is significantly different (Figure 9A and B).

**Figure 8.**
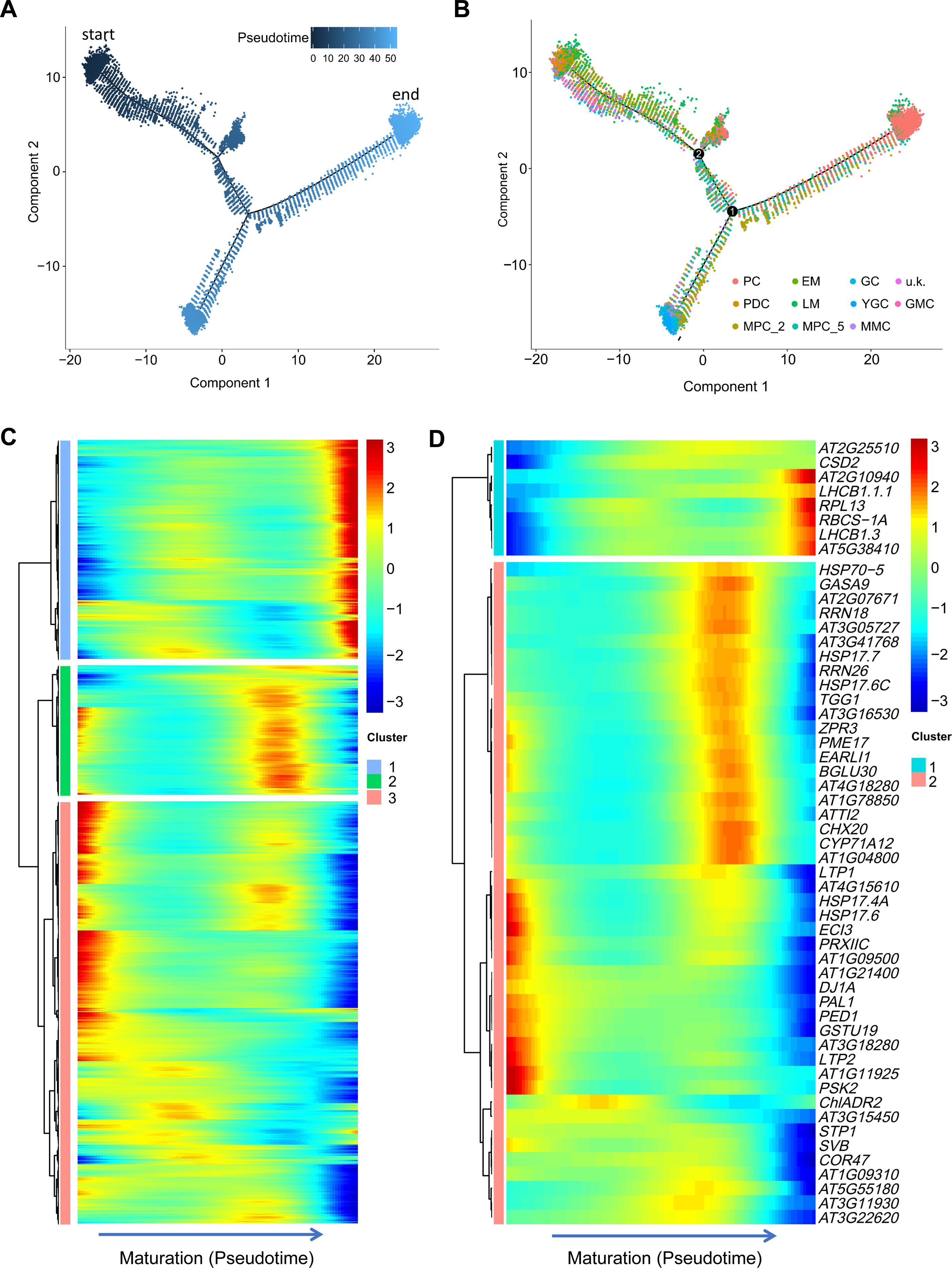
Pseudotime analysis of clusters and the selected marker genes. (**A**) Distribution of cells of each cluster on the pseudotime trajectory. (**B**) Pseudotime trajectory of single-cell transcriptomics data colored according to the cluster labels. Most cells were distributed along main stem, although two small branches were detected near the main path. (**C**) Clustering of all genes during pseudotime progression. (**D**) Clustering and expression kinetics of representative genes along pseudotime progression of stomatal lineage cells.

**Figure 9.**
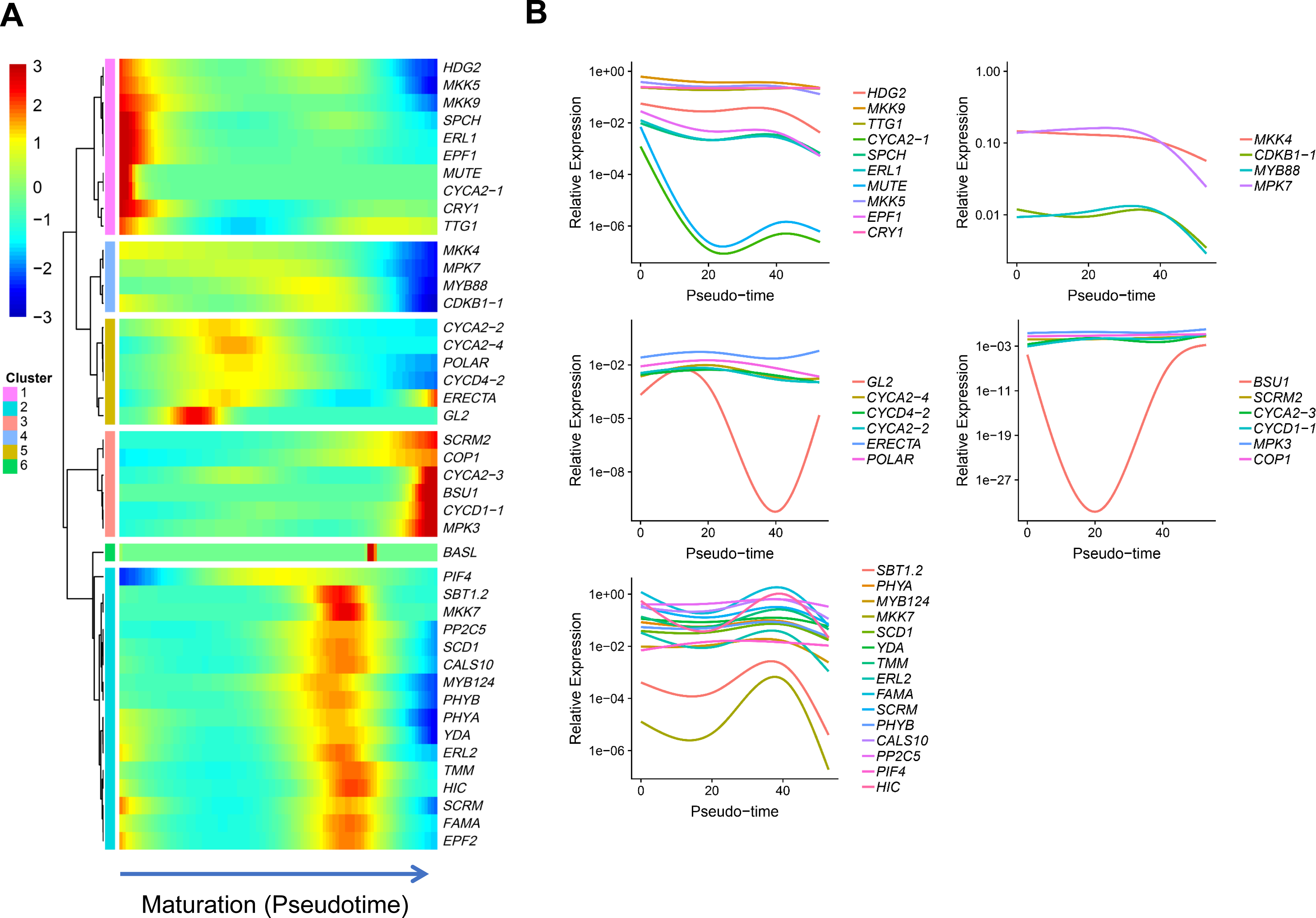
Pseudo-time analysis of known marker genes. (**A**) Clustering of representative genes along pseudo-time progression of stomatal lineage cells. (**B**) Gene expression kinetics along pseudo-time progression of representative genes.

## DISCUSSION

Stomatal lineage cell fate decisions are traceable, irreversible, and produce well-known differentiated cell types. We were able to investigate the interplay of multiple fate-specific genetic programs and the effects of external environmental factors on the fate decision of different cell types at the single cell level. A combination of known marker genes and GO analysis enabled us to reliably classify and define cell types (Figure 1, Figure S3). Transcripts of some marker genes (*SPCH*, *MUTE* and *FAMA*) are enriched in specific types of stomatal lineage cells (Figure S3), while those of other marker genes (*CRY1*, *PP2C*, *BCA4* and *CALS10*) are not (Figure S3). Furthermore, a series of new marker genes were identified in different cell types (Figure 2). We also analyzed the effects of some marker genes on stomatal lineage cell development by checking the stomatal developmental patterns in the cotyledons of the corresponding mutants (Figures 3, 6, and 7).

### Determination of cell types and marker genes

The stomatal lineage cells can be classified into seven different types including PDC, MMC, EM, LM, GMC, YGC and GC based on their developmental stages. To determine the cell type we analyzed the feature plot of the selected marker genes in specific cell types (Figure S3B). We used more than one marker genes for identifying one cell type because marker genes are often expressed in more than one cell type and to different levels (Figure 1). In the case of PDC, no known marker genes can be used, but some marker genes are possibly expressed in PDC before entry into the stomatal lineage cell pathway, such as *EFP2*, *TMM*, *BASL*, *SPCH* although their expression levels are less than in MMC. The MMC, EM, LM, YGC and GC can be clearly identified based on the existence of specific marker genes in each cluster. PC contains chloroplasts and is difficult to distinguish from MPC (Figure 1C). Auxin is required for the formation of the interdigitated cell pattern in leaf pavement cells through the coordination of the two mutually exclusive ROP2 and ROP6 pathways (Xu et al., 2011). The auxin-activated ROP2 pathway is essential for PIN1 polar localization at the lobe apex by inhibiting its internalization (Xu et al., 2010). Therefore, we used ROP2 and ROP6 as marker genes to identify PC. Based on these considerations we propose that cluster 0 represents PC. We could not determine the cell type of cluster 9 with known marker genes (Figure 1C). Interestingly, we found that one marker gene, *BAG6*, may be involved in regulating the biogenesis and distribution of stomata (Figure S4). However, *bZIP6*, another marker gene of cluster 9, is specifically expressed in the two pericycles in the phloem pole starting from the early root elongation zone, suggesting that cluster 9 does not belong to epidermal cells. We selected the top 10 marker genes in each cluster as representative marker genes for the different cell types (Figure 2). Analysis of the marker genes in GC indicated that some of them are involved in regulating the development and function of stomata (Figure 2A), suggesting that the marker genes identified can be used for determining the cell type.

### Potential factors that regulate the fate of stomatal lineage cells

The stomatal lineage cell development is regulated by many important factors, such as light, temperature, metabolism, and phytohormones (Pillitteri and Dong, 2013). In the past, most studies using whole plant cells, could not clearly distinguish the special functions of these factors in the different cell types. For instance, light- and hormone-signaling can affect the entire process of stomatal lineage cell development (Pillitteri and Dong, 2013). Based on scRNA-seq, combined with GO analysis, we were able to identify the potential genes that regulate stomatal lineage cell development. For example, GO heatmap analysis revealed that genes preferentially expressed in YGC and GC are mainly involved in the response to oxidative stress, abscisic acid, osmotic stress, and vacuolar activity (Figure 4). The GO heatmap analysis also showed that GO terms enriched in MMC, EM, LM, and GMC are relatively similar (Figure 4A), suggesting there are intense interactions in gene expression and cell functions among these cells. GO terms enriched in LM and GMC are involved in regulating the respiratory chain (Figure 4B), suggesting that the differentiation from LM to GMC is an energy-intensive process. It should be noted that genes highly expressed in MMC are involved in the response to bacterial infection and in the MAPK signaling pathway (Figure 4B). Studies have shown that bacterial infection can generate a systemic signal that is translocated from the mature infected leaves to the developing leaves in the apical meristem, where it reduces stomatal density by increasing epidermal cell expansion in the newly developing leaves (Dutton et al., 2019). After infection fewer epidermal cells enter the stomatal lineage during the early stages of leaf development (Dutton et al., 2019). Taken together our results indicate that, genes expressed in MMC are required for suppressing the biogenesis of stomatal cells in response to bacterial infection. Interestingly, GO analysis revealed that genes expressed in PDCs and MMCs are involved in the regulation of protein transport, vesicle-mediated transport and in membrane protein complexes (Figure 4A).

Analysis of the new identified marker genes revealed that expression of *ATML1* and *PDF1* was specifically enhanced in MMC. *ATML1* and *PDF1* show high co-expression with *MUTE* and *SPCH* (Figure 2A and Figure S3A). Further analysis revealed that *atml1-2* and *atml1-3* are deficient in the development of stomatal lineage cells (Figure 3). In addition, the expression levels of both *SPCH* and *MUTE* were decreased in *atml1-2* and *atml1-3*, compared with WT (Figure 3E). *ATML1* encodes a homeobox protein similar to *GL2* and is expressed in both the apical and basal daughter cells of the zygote as well as in its progeny (Peterson et al., 2013). Expression of *ATML1* starts at the two-cell stage of embryo development and is later restricted to the outermost epidermal cell layer (Iida et al., 2019). The *ATML1* promoter is highly modular with each of its domains contributing to specific features of the spatial and temporal expression of the gene (Takada et al., 2013). Double mutant analysis with *pdf2*, another L1-specific gene, suggests that their functions are partially redundant, since the loss of both genes results in abnormal shoot development (Ogawa et al., 2015). Over-expression of *ATML1* can induce the formation of stomata-like structures in the inner cells of the cotyledons in independent lines (Peterson et al., 2013). Therefore, taken together, our results suggest that ATML1 can regulate the development of stomatal lineage cells by modulating the expression of both *SPCH* and *MUTE*.

### Involvement of a TF regulatory network in regulating stomatal lineage cell development

It is well known that bHLH TFs play important roles in regulating stomatal lineage cell development (MacAlister et al., 2007; Pillitteri et al., 2007). Recently, additional new TFs that are involved in regulating the stomatal lineage cell development have been identified, for example PIF4, MYB88, HDG2, GL2 (Casson et al., 2009; Pillitteri and Dong, 2013). To identify new TFs that regulate stomatal lineage cell development in special cell types, we analyzed potential TFs expressed in different stomatal lineage cells. Analysis of the network of TFs indicated that PIF5, WRKY33, BPC1, and BPC6 may act as the core TFs that regulate the differentiation from PDC to GMC (Figure 5C and D). The BPC gene family has seven members in Arabidopsis (Monfared et al., 2011). BPC belongs to GAGA binding proteins (GBPs), which bind GA-rich elements (Biggin and Tjian, 1988; Kooiker et al., 2005; Monfared et al., 2011). These GBPs are involved in regulating gene expression by interacting with chromatin remodeling complexes like NURF and FACT (Lehmann, 2004). Expression of *BPC* genes occurs widely, but to different extents, in various organs. These genes play important roles in regulating the vegetative and reproductive development (Kooiker et al., 2005; Monfared et al., 2011; Simonini et al., 2012; Simonini and Kater, 2014; Mu et al., 2017; Shanks et al., 2018). Our results indicate that besides the core TFs (e.g. SPCH, MUTE, FAMA, etc), a TF network comprising WRKY33, BPC1/6, and PIF5 is required for modulating the development of stomatal cells. Further analysis revealed that *wrky33* and the *bpc1 bpc2 bpc4 bpc6* quadruple mutant are deficient in different stomatal lineage cells (Figure 6 and Figure 7). The expression levels of *SCRM* and *SCRM2* in the *bpc1 bpc2 bpc3 bpc4 bpc6 bpc7* sextuple mutant were less than in WT (Figure 7). These results suggest that WRKY33, BPCs, and PIF5 act as the core TFs that regulate the differentiation from PDC to GMC.

### The developmental trajectory of stomatal lineage cells

To dissect the temporal and spatial distribution of stomatal lineage cells we performed a pseudo-time analysis on scRNAseq data (Figure 8A and B). Pseudo-time patterns of all genes can be divided into three clusters (Figure 8C). Further analysis of the top 5 selected marker genes in each cluster indicated that these marker genes can be grouped into two clusters (Figure 8D). The first cluster mainly contains the markers from MPC (Figure 8D). The second cluster can be divided into three sub-clusters: the first sub-cluster includes the marker genes from GMC, YGC, GC and MPC; the second sub-cluster contains the marker genes from PDC and LM; and the third sub-cluster includes the marker genes from MMC and EM (Figure 8D). As typical marker genes, *SPCH* and *MUTE* were found to be co-expressed with *EPF1*, *MKK5* and *MKK9* (Figure 9A and B), suggesting that EPF1-MKK9/5 dependent signaling can influence both the expression and the function of SPCH and MUTE. More interestingly, light-signal receptor genes and stomatal lineage marker genes show strong co-expression patterns (Figure 9A). In response to changes in light quality, different light signal receptors rely on downstream COP1-YDA-MAPK signaling pathways to regulate different stages of stomatal lineage development (Kang et al., 2009). Our results show that *SCRM2* and *COP1* exhibit very similar pseudo-time curves, while *SCRM* has the same pseudo-time curves as *PHYA* and *PHYB* (Figure 9A and B). Unlike *PHYA* and *PHYB*, the pseudo-time curves of the blue receptor *CRY1* is significantly different from those of *PHYA* and *PHYB* (Figure 9A and B). Surprisingly, although *EPF1* and *EPF2* play very similar roles in regulating stomatal lineage cell development, they exhibit distinct expression patterns (Figure 9A and B). It has been reported that EPF2 activates ER signaling, leading to subsequent MAPK activation and inhibition of stomatal lineage cell development, while EPFL9 prevents signal transduction of MPK3 and MPK6 (Lee et al., 2015). In the pseudo-time course, however, we observed that *MPK3* has the exact opposite expression pattern compared with *EPF1* (Figure 9A and B).

The transition to GMC is coordinated through cell-cycle controls and is promoted by MUTE (Han et al., 2018), while FAMA and FLP/MYB88 act in parallel to antagonize GMC transition (Lai et al., 2005). The canonical G1 and G1/S-regulating CYCD family member *CYCD5;1* is a MUTE target, implying that it may promote symmetric cell division (SCD) commitment in a MUTE-dependent manner (Han et al., 2018). Interestingly, *CYCD7;1* is not regulated by MUTE, although *CYCD7;1* is specifically expressed in stomatal lineage cells (Adrian et al., 2015). In addition to *CYCD5;1*, our results reveal that expression of *CYCA2;1* is highly similar to that of *MUTE* and *SPCH* (Figures 9A and B). However, the expression of *CYCA2;1* is restricted to the vascular tissues of leaves (Vanneste et al., 2011). The co-expression of *MUTE* and *CYCA2;1* imply that the expression of these two genes can be induced at a similar development time, but the expression of *CYCA2;1* may not be regulated by MUTE.

## METHODS

### Screening and Verification of Mutants

T-DNA insertion mutants were obtained from the Arabidopsis Biological Resource Center (ABRC) (Supplemental **Table 3**). Mutant lines homozygous for the T-DNA insertion were identified by PCR analysis using gene-specific and T-DNA-specific primers (Supplemental **Table 4**). In addition, we also generated the transgenic lines of *BPC6-GFP*.

### Constructs for plant transformation

To generate the pB7WGF2-*BPC6* constructs, the full-length cDNA of *BPC6* was PCR-amplified using the primer pairs as described in Supplementary **Table 2**. Then the PCR products were purified, and first cloned into pDNOR201 by BP Clonase reactions (GATEWAY Cloning; Invitrogen) according to the manufacturer’s instructions to generate the pDNOR-*BPC6*. The resulting plasmids were recombined into pB7WGF2 using LR Clonase reactions (GATEWAY Cloning; Invitrogen) to generate the final constructs.

## PLANT TRANSFORMATION

The pB7WGF2-*BPC6* constructs were transformed into *Agrobacterium tumefaciens* strain GV3105 *via* electroporation. Then the *Agrobacterium tumefaciens* that contained the constructs of pB7WGF2-*BPC6* was introduced into WT. The resulting T1 transgenic plants of pB7WGF2-*BPC6* were selected by BASTA as described by Sun et al (Sun et al., 2016). Homozygous transgenic plants were used in all experiments.

### Cotyledon Collection and Protoplast Preparation

We isolated protoplasts from cotyledons of five-day-old *Arabidopsis* seedlings as described by Yoo, et al., (2007)(Yoo et al., 2007) with slight modifications to adjust to the cotyledon tissue. Briefly, the cotyledons were harvested from seedlings submerged in a solution (0.5 mM CaCl_2_, 0.5 mM MgCl_2_, 5 mM MES, 1.5% Cellulase RS, 0.03% Pectolyase Y23, 0.25% BSA, actinomycin D [33 mg/L], and cordycepin [100mg/L], pH 5.5) by vacuum infiltration for 10 min. The samples were then incubated for 4 hours to isolate protoplasts. Afterwards, the isolated cells were washed three times with 8% mannitol buffer to remove Mg^2+^. Cells were then filtered with a 40 µm cell strainer. Cell activity was detected by trypan blue staining and cell concentration was measured with a hemocytometer.

### Single-cell RNA-seq Library Preparation

We prepared single-cell RNA-seq libraries with Chromium Single Cell 3ʹ Gel Beads-in-emulsion (GEM) Library & Gel Bead Kit v3 according to the user manual supplied by the kit. In brief, GEMs were generated and barcoded, followed by post GEM-RT cleanup and cDNA Amplification, and finally 3ʹ Gene Expression Library Construction. In the first step cells were diluted so that the majority (∼90-99%) of GEMs contained no cells, while the remainder mostly contained a single cell. The Gel Beads were then dissolved, primers were released, and any co-partitioned cell was lysed in order to generate full-length cDNA from poly-adenylated mRNA. Subsequently, the first-strand cDNA from the post GEM-RT reaction mixture was purified with SILANE magnetic beads. After purification, the barcoded full-length cDNA was amplified via PCR to generate sufficient amounts for library construction. In the third step, enzymatic fragmentation and size selection were used to optimize the cDNA amplicon size. In addition, the TruSeq Read 1 (read 1 primer sequence) was added to the molecules during GEM incubation. P5, P7, a sample index, and TruSeq Read 2 (read 2 primer sequence) were added via End Repair. This was followed by A-tailing, adaptor ligation, and PCR. The final libraries contained the P5 and P7 primers used in Illumina bridge amplification.

### Single-cell RNA-seq Data Preprocessing

The Cell Ranger pipeline (version 3.0.0) provided by 10× Genomics was used to demultiplex cellular barcodes and map reads to the TAIR10 reference genome. Transcript quantifications were determined via the STAR aligner. We processed the unique molecular identifier (UMI) count matrix using the R package Seurat (version 2.3.4). To remove low quality cells and likely multiple captures, we further applied criteria to filter out cells with UMI/gene numbers outside the limit of the mean value +/- 2 standard deviations, assuming a Gaussian distribution of each cell’s UMI/gene numbers. Following visual inspection of the distribution of cells by the fraction of chloroplast genes expressed, we further discarded low-quality cells where >40% of the counts belonged to chloroplast genes. After applying these quality control (QC) criteria, 12,844 single cells and 32,833 genes in total remained and were included in the downstream analyses. Library size normalization was performed in Seurat on the filtered matrix to obtain normalized counts.

Genes with the highest variable expression amongst single cells were identified using the method described in Macosko et al (Macosko et al., 2015). Briefly, the average expression and dispersion were calculated for all genes, which were subsequently placed into 11 bins based on expression. Principal component analysis (PCA) was performed to reduce the dimensionality on the log transformed gene-barcode matrices of the most variable genes. Cells were clustered via a graph-based approach and visualized in 2-dimensions using tSNE. A likelihood ratio test, which simultaneously tests for changes in mean expression and percentage of cells expressing a gene, was used to identify significantly differentially expressed genes (DEGs) between clusters. We also performed tSNE analyses and identified the DEGs between clusters for the mesophyll and stomatal lineage cell populations.

Pseudotime trajectory analysis of single cell transcriptomes was conducted using Monocle 2 (Trapnell et al., 2014). Genes with the most highly variable expression were used for clustering the cells. Gene expression was then plotted as a function of pseudo-time in Monocle 2 to track changes across pseudo-time. We also plotted TFs and marker genes along the inferred developmental pseudo-time. The regulation networks for the TFs and target genes were plotted by Cytoscape according to the PlantTFDB database.

### Microscopy

The cotyledons were observed 5 d after germination. The samples were harvested and placed in 70% ethanol, cleared overnight at room temperature, and then stored in Hoyer’s Solution. Images of stomata were obtained from samples stored in Hoyer’s Solution and visualized using differential interference contrast microscopy with a Leica DMi8 microscope. A Nikon D-ECLIPSE C1 laser confocal scanning microscope was used for green fluorescent protein (GFP) fluorescence images.

### Gene Ontology (GO) Enrichment Analysis

The enrichment of gene ontology (GO) terms and pathways for the DEGs were analyzed using Metascape (http://metascape.org/) (Zhou et al., 2019).

### Accession Numbers

Sequence data from this study can be found in the Arabidopsis Genome Initiative data library under the following accession numbers: *WRKY33* (AT1G07890), *BPC1* (AT2G01930), *BPC2* (AT1G14685), *BPC3* (AT1G68120), *BPC4* (AT2G21240), *BPC6* (AT5G42520), *BPC7* (AT2G35550), *ATML1* (AT4G21750), *BAG6* (AT2G46240), *bZIP6* (AT2G22850). Single cell RNA sequence data are available at the https://dataview.ncbi.nlm.nih.gov/?search=SUB6947465 (https://www.ncbi.nlm.nih.gov).

## ACKNOWLEDGEMENTS

We are grateful to ABRC for the Arabidopsis seeds. This research was supported by the National Natural Science Foundation of China (31670233) and the Program of Introducing Talents of Discipline to Universities. We thank Prof. Charles Gasser for providing the *bpc double, triple, quadruple, and sextuple* mutant seeds.

## AUTHOR CONTRIBUTIONS

XS designed the study; ZL, YZ, JG, ZZ, JL, ZT, JW, RW, BZ, WL, TL, YH and YH performed the research; RJ, YM and XS analyzed the data; XS and RJ wrote the paper. All authors discussed the results and made comments on the manuscript.

## REFERENCES

Adrian, J., Chang, J., Ballenger, C.E., Bargmann, B.O.R., Alassimone, J., Davies, K.A., Lau, O.S., Matos, J.L., Hachez, C., Lanctot, A., Vaten, A., Birnbaum, K.D., and Bergmann, D.C. (2015). Transcriptome Dynamics of the Stomatal Lineage: Birth, Amplification, and Termination of a Self-Renewing Population. Developmental Cell 33, 107–118.

Amsbury, S., Hunt, L., Elhaddad, N., Baillie, A., Lundgren, M., Verhertbruggen, Y., Scheller, H.V., Knox, J.P., Fleming, A.J., and Gray, J.E. (2016). Stomatal Function Requires Pectin De-methyl-esterification of the Guard Cell Wall. Current Biology 26, 2899–2906.

Barton, K.A., Schattat, M.H., Jakob, T., Hause, G., Wilhelm, C., McKenna, J.F., Mathe, C., Runions, J., Van Damme, D., and Mathur, J. (2016). Epidermal Pavement Cells of Arabidopsis Have Chloroplasts. Plant Physiol 171, 723–726.

Bergmann, D.C., and Sack, F.D. (2007). Stomatal development. Annu Rev Plant Biol 58, 163–181.

Bergmann, D.C., Lukowitz, W., and Somerville, C.R. (2004). Stomatal development and pattern controlled by a MAPKK kinase. Science 304, 1494–1497.

Biggin, M.D., and Tjian, R. (1988). Transcription factors that activate the Ultrabithorax promoter in developmentally staged extracts. Cell 53, 699–711.

Casson, S.A., Franklin, K.A., Gray, J.E., Grierson, C.S., Whitelam, G.C., and Hetherington, A.M. (2009). phytochrome B and PIF4 Regulate Stomatal Development in Response to Light Quantity. Current Biology 19, 229–234.

Chanroj, S., Padmanaban, S., Czerny, D.D., Jauh, G.Y., and Sze, H. (2013). K+ transporter AtCHX17 with its hydrophilic C tail localizes to membranes of the secretory/endocytic system: role in reproduction and seed set. Mol Plant 6, 1226–1246.

Chen, L., Guan, L.P., Qian, P.P., Xu, F., Wu, Z.L., Wu, Y.J., He, K., Gou, X.P., Li, J., and Hou, S.W. (2016). NRPB3, the third largest subunit of RNA polymerase II, is essential for stomatal patterning and differentiation in Arabidopsis. Development 143, 1600–1611.

Dong, J., MacAlister, C.A., and Bergmann, D.C. (2009). BASL controls asymmetric cell division in Arabidopsis. Cell 137, 1320–1330.

Dutton, C., Horak, H., Hepworth, C., Mitchell, A., Ton, J., Hunt, L., and Gray, J.E. (2019). Bacterial infection systemically suppresses stomatal density. Plant Cell Environ 42, 2411–2421.

Engineer, C.B., Ghassemian, M., Anderson, J.C., Peck, S.C., Hu, H.H., and Schroeder, J.I. (2015). Carbonic anhydrases, EPF2 and a novel protease mediate CO2 control of stomatal development (vol 513, pg 246, 2014). Nature 526, 458–458.

Fu, C., Hou, Y., Ge, J., Zhang, L., Liu, X., Huo, P., and Liu, J. (2019). Increased fes1a thermotolerance is induced by BAG6 knockout. Plant Mol Biol 100, 73–82.

Fu, Y., Gu, Y., Zheng, Z., Wasteneys, G., and Yang, Z. (2005). Arabidopsis interdigitating cell growth requires two antagonistic pathways with opposing action on cell morphogenesis. Cell 120, 687–700.

Geisler, M., Nadeau, J., and Sack, F.D. (2000). Oriented asymmetric divisions that generate the stomatal spacing pattern in Arabidopsis are disrupted by the too many mouths mutation. Plant Cell 12, 2075–2086.

Gray, J. (2005). Guard cells: transcription factors regulate stomatal movements. Curr Biol 15, R593–595.

Han, S.K., and Torii, K.U. (2016). Lineage-specific stem cells, signals and asymmetries during stomatal development. Development 143, 1259–1270.

Han, S.K., Qi, X., Sugihara, K., Dang, J.H., Endo, T.A., Miller, K.L., Kim, E.D., Miura, T., and Torii, K.U. (2018). MUTE Directly Orchestrates Cell-State Switch and the Single Symmetric Division to Create Stomata. Dev Cell 45, 303–315 e305.

Hara, K., Kajita, R., Torii, K.U., Bergmann, D.C., and Kakimoto, T. (2007). The secretory peptide gene EPF1 enforces the stomatal one-cell-spacing rule. Gene Dev 21, 1720–1725.

Hronkova, M., Wiesnerova, D., Simkova, M., Skupa, P., Dewitte, W., Vrablova, M., Zazimalova, E., and Santrucek, J. (2015). Light-induced STOMAGEN-mediated stomatal development in Arabidopsis leaves. Journal of Experimental Botany 66, 4621–4630.

Huang, S., Waadt, R., Nuhkat, M., Kollist, H., Hedrich, R., and Roelfsema, M.R.G. (2019). Ca(2+) signals in guard cells enhance the efficiency by which ABA triggers stomatal closure. New Phytol.

Hunt, L., and Gray, J.E. (2009a). The signaling peptide EPF2 controls asymmetric cell divisions during stomatal development. Curr Biol 19, 864–869.

Hunt, L., and Gray, J.E. (2009b). The Signaling Peptide EPF2 Controls Asymmetric Cell Divisions during Stomatal Development. Current Biology 19, 864–869.

Iida, H., Yoshida, A., and Takada, S. (2019). ATML1 activity is restricted to the outermost cells of the embryo through post-transcriptional repressions. Development 146.

Kabbage, M., and Dickman, M.B. (2008). The BAG proteins: a ubiquitous family of chaperone regulators. Cell Mol Life Sci 65, 1390–1402.

Kanaoka, M.M., Pillitteri, L.J., Fujii, H., Yoshida, Y., Bogenschutz, N.L., Takabayashi, J., Zhu, J.K., and Torii, K.U. (2008). SCREAM/ICE1 and SCREAM2 specify three cell-state transitional steps leading to Arabidopsis stomatal differentiation. Plant Cell 20, 1775–1785.

Kang, C.Y., Lian, H.L., Wang, F.F., Huang, J.R., and Yang, H.Q. (2009). Cryptochromes, Phytochromes, and COP1 Regulate Light-Controlled Stomatal Development in Arabidopsis. Plant Cell 21, 2624–2641.

Kim, T.W., Michniewicz, M., Bergmann, D.C., and Wang, Z.Y. (2012). Brassinosteroid regulates stomatal development by GSK3-mediated inhibition of a MAPK pathway. Nature 482, 419–U1526.

Kooiker, M., Airoldi, C.A., Losa, A., Manzotti, P.S., Finzi, L., Kater, M.M., and Colombo, L. (2005). BASIC PENTACYSTEINE1, a GA binding protein that induces conformational changes in the regulatory region of the homeotic Arabidopsis gene SEEDSTICK. Plant Cell 17, 722–729.

Lai, L.B., Nadeau, J.A., Lucas, J., Lee, E.K., Nakagawa, T., Zhao, L.M., Geisler, M., and Sack, F.D. (2005). The Arabidopsis R2R3 MYB proteins FOUR LIPS and MYB88 restrict divisions late in the stomatal cell lineage. Plant Cell 17, 2754–2767.

Lampard, G.R., MacAlister, C.A., and Bergmann, D.C. (2008). Arabidopsis Stomatal Initiation Is Controlled by MAPK-Mediated Regulation of the bHLH SPEECHLESS. Science 322, 1113–1116.

Lampard, G.R., Lukowitz, W., Ellis, B.E., and Bergmann, D.C. (2009). Novel and Expanded Roles for MAPK Signaling in Arabidopsis Stomatal Cell Fate Revealed by Cell Type-Specific Manipulations. Plant Cell 21, 3506–3517.

Lawson, T. (2009). Guard cell photosynthesis and stomatal function. New Phytol 181, 13–34.

Lee, J.S., Hnilova, M., Maes, M., Lin, Y.C.L., Putarjunan, A., Han, S.K., Avila, J., and Torii, K.U. (2015). Competitive binding of antagonistic peptides fine-tunes stomatal patterning. Nature 522, 435-+.

Lee, J.S., Kuroha, T., Hnilova, M., Khatayevich, D., Kanaoka, M.M., McAbee, J.M., Sarikaya, M., Tamerler, C., and Torii, K.U. (2012). Direct interaction of ligand-receptor pairs specifying stomatal patterning. Gene Dev 26, 126–136.

Lee, J.Y., Colinas, J., Wang, J.Y., Mace, D., Ohler, U., and Benfey, P.N. (2006). Transcriptional and posttranscriptional regulation of transcription factor expression in Arabidopsis roots. Proc Natl Acad Sci U S A 103, 6055–6060.

Lehmann, M. (2004). Anything else but GAGA: a nonhistone protein complex reshapes chromatin structure. Trends in genetics : TIG 20, 15–22.

Li, J., Brader, G., and Palva, E.T. (2008). Kunitz trypsin inhibitor: an antagonist of cell death triggered by phytopathogens and fumonisin b1 in Arabidopsis. Mol Plant 1, 482–495.

Li, Y., Kabbage, M., Liu, W., and Dickman, M.B. (2016). Aspartyl Protease-Mediated Cleavage of BAG6 Is Necessary for Autophagy and Fungal Resistance in Plants. Plant Cell 28, 233–247.

Liang, H., Zhang, Y., Martinez, P., Rasmussen, C.G., Xu, T., and Yang, Z. (2018). The Microtubule-Associated Protein IQ67 DOMAIN5 Modulates Microtubule Dynamics and Pavement Cell Shape. Plant Physiol 177, 1555–1568.

MacAlister, C.A., Ohashi-Ito, K., and Bergmann, D.C. (2007). Transcription factor control of asymmetric cell divisions that establish the stomatal lineage. Nature 445, 537–540.

Macosko, E.Z., Basu, A., Satija, R., Nemesh, J., Shekhar, K., Goldman, M., Tirosh, I., Bialas, A.R., Kamitaki, N., Martersteck, E.M., Trombetta, J.J., Weitz, D.A., Sanes, J.R., Shalek, A.K., Regev, A., and McCarroll, S.A. (2015). Highly Parallel Genome-wide Expression Profiling of Individual Cells Using Nanoliter Droplets. Cell 161, 1202–1214.

Maresova, L., and Sychrova, H. (2006). Arabidopsis thaliana CHX17 gene complements the kha1 deletion phenotypes in Saccharomyces cerevisiae. Yeast 23, 1167–1171.

Melotto, M., Underwood, W., Koczan, J., Nomura, K., and He, S.Y. (2006). Plant stomata function in innate immunity against bacterial invasion. Cell 126, 969–980.

Monfared, M.M., Simon, M.K., Meister, R.J., Roig-Villanova, I., Kooiker, M., Colombo, L., Fletcher, J.C., and Gasser, C.S. (2011). Overlapping and antagonistic activities of BASIC PENTACYSTEINE genes affect a range of developmental processes in Arabidopsis. Plant J 66, 1020–1031.

Mu, Y., Zou, M., Sun, X., He, B., Xu, X., Liu, Y., Zhang, L., and Chi, W. (2017). BASIC PENTACYSTEINE Proteins Repress ABSCISIC ACID INSENSITIVE4 Expression via Direct Recruitment of the Polycomb-Repressive Complex 2 in Arabidopsis Root Development. Plant Cell Physiol 58, 607–621.

Mucha, S., Heinzlmeir, S., Kriechbaumer, V., Strickland, B., Kirchhelle, C., Choudhary, M., Kowalski, N., Eichmann, R., Huckelhoven, R., Grill, E., Kuster, B., and Glawischnig, E. (2019). The Formation of a Camalexin Biosynthetic Metabolon. Plant Cell 31, 2697–2710.

Muller, T.M., Bottcher, C., Morbitzer, R., Gotz, C.C., Lehmann, J., Lahaye, T., and Glawischnig, E. (2015). TRANSCRIPTION ACTIVATOR-LIKE EFFECTOR NUCLEASE-Mediated Generation and Metabolic Analysis of Camalexin-Deficient cyp71a12 cyp71a13 Double Knockout Lines. Plant Physiol 168, 849–858.

Nadeau, J.A., and Sack, F.D. (2002). Control of stomatal distribution on the Arabidopsis leaf surface. Science 296, 1697–1700.

Nafisi, M., Goregaoker, S., Botanga, C.J., Glawischnig, E., Olsen, C.E., Halkier, B.A., and Glazebrook, J. (2007). Arabidopsis cytochrome P450 monooxygenase 71A13 catalyzes the conversion of indole-3-acetaldoxime in camalexin synthesis. Plant Cell 19, 2039–2052.

Niu, M.L., Huang, Y., Sun, S.T., Sun, J.Y., Cao, H.S., Shabala, S., and Bie, Z.L. (2018). Root respiratory burst oxidase homologue-dependent H2O2 production confers salt tolerance on a grafted cucumber by controlling Na+ exclusion and stomatal closure. Journal of Experimental Botany 69, 3465–3476.

Ogawa, E., Yamada, Y., Sezaki, N., Kosaka, S., Kondo, H., Kamata, N., Abe, M., Komeda, Y., and Takahashi, T. (2015). ATML1 and PDF2 Play a Redundant and Essential Role in Arabidopsis Embryo Development. Plant Cell Physiol 56, 1183–1192.

Ohashi-Ito, K., and Bergmann, D.C. (2006). Arabidopsis FAMA controls the final proliferation/differentiation switch during stomatal development. Plant Cell 18, 2493–2505.

Peterson, K.M., Shyu, C., Burr, C.A., Horst, R.J., Kanaoka, M.M., Omae, M., Sato, Y., and Torii, K.U. (2013). Arabidopsis homeodomain-leucine zipper IV proteins promote stomatal development and ectopically induce stomata beyond the epidermis. Development 140, 1924–1935.

Pillitteri, L.J., and Torii, K.U. (2012). Mechanisms of Stomatal Development. Annu Rev Plant Biol 63, 591–614.

Pillitteri, L.J., and Dong, J. (2013). Stomatal development in Arabidopsis. Arabidopsis Book 11, e0162.

Pillitteri, L.J., Sloan, D.B., Bogenschutz, N.L., and Torii, K.U. (2007). Termination of asymmetric cell division and differentiation of stomata. Nature 445, 501–505.

Pilot, G., Lacombe, B., Gaymard, F., Cherel, I., Boucherez, J., Thibaud, J.B., and Sentenac, H. (2001). Guard cell inward K+ channel activity in arabidopsis involves expression of the twin channel subunits KAT1 and KAT2. J Biol Chem 276, 3215–3221.

Pitzschke, A., Xue, H., Persak, H., Datta, S., and Seifert, G.J. (2016). Post-Translational Modification and Secretion of Azelaic Acid Induced 1 (AZI1), a Hybrid Proline-Rich Protein from Arabidopsis. Int J Mol Sci 17.

Rajniak, J., Barco, B., Clay, N.K., and Sattely, E.S. (2015). A new cyanogenic metabolite in Arabidopsis required for inducible pathogen defence. Nature 525, 376–379.

Rudall, P.J., Hilton, J., and Bateman, R.M. (2013). Several developmental and morphogenetic factors govern the evolution of stomatal patterning in land plants. New Phytologist 200, 598–614.

Shanks, C.M., Hecker, A., Cheng, C.Y., Brand, L., Collani, S., Schmid, M., Schaller, G.E., Wanke, D., Harter, K., and Kieber, J.J. (2018). Role of BASIC PENTACYSTEINE transcription factors in a subset of cytokinin signaling responses. Plant J 95, 458–473.

Shpak, E.D., McAbee, J.M., Pillitteri, L.J., and Torii, K.U. (2005). Stomatal patterning and differentiation by synergistic interactions of receptor kinases. Science 309, 290–293.

Simonini, S., and Kater, M.M. (2014). Class I BASIC PENTACYSTEINE factors regulate HOMEOBOX genes involved in meristem size maintenance. J Exp Bot 65, 1455–1465.

Simonini, S., Roig-Villanova, I., Gregis, V., Colombo, B., Colombo, L., and Kater, M.M. (2012). Basic pentacysteine proteins mediate MADS domain complex binding to the DNA for tissue-specific expression of target genes in Arabidopsis. Plant Cell 24, 4163–4172.

Song, Y., Miao, Y., and Song, C.P. (2014). Behind the scenes: the roles of reactive oxygen species in guard cells. New Phytol 201, 1121–1140.

Sugano, S.S., Shimada, T., Imai, Y., Okawa, K., Tamai, A., Mori, M., and Hara-Nishimura, I. (2010). Stomagen positively regulates stomatal density in Arabidopsis. Nature 463, 241–U130.

Sun, X., Xu, D., Liu, Z., Kleine, T., and Leister, D. (2016). Functional relationship between mTERF4 and GUN1 in retrograde signaling. J Exp Bot 67, 3909–3924.

Szyroki, A., Ivashikina, N., Dietrich, P., Roelfsema, M.R.G., Ache, P., Reintanz, B., Deeken, R., Godde, M., Felle, H., Steinmeyer, R., Palme, K., and Hedrich, R. (2001). KAT1 is not essential for stomatal opening. P Natl Acad Sci USA 98, 2917–2921.

Takada, S., Takada, N., and Yoshida, A. (2013). ATML1 promotes epidermal cell differentiation in Arabidopsis shoots. Development 140, 1919–1923.

Trapnell, C., Cacchiarelli, D., Grimsby, J., Pokharel, P., Li, S., Morse, M., Lennon, N.J., Livak, K.J., Mikkelsen, T.S., and Rinn, J.L. (2014). The dynamics and regulators of cell fate decisions are revealed by pseudotemporal ordering of single cells. Nat Biotechnol 32, 381–386.

Underwood, W., Melotto, M., and He, S.Y. (2007). Role of plant stomata in bacterial invasion. Cell Microbiol 9, 1621–1629.

Vanneste, S., Coppens, F., Lee, E., Donner, T.J., Xie, Z., Van Isterdael, G., Dhondt, S., De Winter, F., De Rybel, B., Vuylsteke, M., De Veylder, L., Friml, J., Inze, D., Grotewold, E., Scarpella, E., Sack, F., Beemster, G.T., and Beeckman, T. (2011). Developmental regulation of CYCA2s contributes to tissue-specific proliferation in Arabidopsis. EMBO J 30, 3430–3441.

von Groll, U., and Altmann, T. (2001). Stomatal cell biology. Current Opinion in Plant Biology 4, 555–560.

Xu, T., Nagawa, S., and Yang, Z. (2011). Uniform auxin triggers the Rho GTPase-dependent formation of interdigitation patterns in pavement cells. Small GTPases 2, 227–232.

Xu, T., Wen, M., Nagawa, S., Fu, Y., Chen, J.G., Wu, M.J., Perrot-Rechenmann, C., Friml, J., Jones, A.M., and Yang, Z. (2010). Cell surface- and rho GTPase-based auxin signaling controls cellular interdigitation in Arabidopsis. Cell 143, 99–110.

Yoo, S.D., Cho, Y.H., and Sheen, J. (2007). Arabidopsis mesophyll protoplasts: a versatile cell system for transient gene expression analysis. Nat Protoc 2, 1565–1572.

Zeng, W., Melotto, M., and He, S.Y. (2010). Plant stomata: a checkpoint of host immunity and pathogen virulence. Curr Opin Biotechnol 21, 599–603.

Zhang, C., Mallery, E.L., and Szymanski, D.B. (2013). ARP2/3 localization in Arabidopsis leaf pavement cells: a diversity of intracellular pools and cytoskeletal interactions. Front Plant Sci 4, 238.

Zhang, D., Tian, C., Yin, K., Wang, W., and Qiu, J.L. (2019). Postinvasive Bacterial Resistance Conferred by Open Stomata in Rice. Mol Plant Microbe Interact 32, 255–266.

Zhang, J.Y., He, S.B., Li, L., and Yang, H.Q. (2014). Auxin inhibits stomatal development through MONOPTEROS repression of a mobile peptide gene STOMAGEN in mesophyll. P Natl Acad Sci USA 111, E3015–E3023.

Zhou, Y., Zhou, B., Pache, L., Chang, M., Khodabakhshi, A.H., Tanaseichuk, O., Benner, C., and Chanda, S.K. (2019). Metascape provides a biologist-oriented resource for the analysis of systems-level datasets. Nat Commun 10, 1523.

